# Tumor suppressor collateral damage screens reveal mRNA homeostasis protein HBS1L as a novel vulnerability in ch9p21 driven FOCAD deleted cancer

**DOI:** 10.1101/2025.02.12.637755

**Authors:** Hongxiang Zhang, Matthew R. Tonini, Lauren Catherine M. Martires, Helena N. Jenkins, Charlotte B. Pratt, Eden N. Gordon, Shanchuan Zhao, Ashley H. Choi, Samuel R. Meier, Tenzing Khendu, Shangtao Liu, Binzhang Shen, Hannah Stowe, Katerina Pashiardis, Xuewen Pan, Madhavi Bandi, Minjie Zhang, Yi Yu, Chengyin Min, Alan Huang, Jannik N. Andersen, Hilary E. Nicholson, Teng Teng

**Author notes:** **Corresponding Author**: Hilary E. Nicholson, 201 Brookline Avenue, Suite 901, Boston, MA 02215. **Conflicts of interest:** All authors were shareholders and/or employees of Tango Therapeutics at the time of their contributions to this body of work.

## Abstract

Chromosomal deletion of tumor suppressor genes often occurs in an imprecise manner, leading to co-deletion of neighboring genes. This collateral damage can create novel dependencies specific to the co-deleted context. One notable example is the dependency on PRMT5 activity in tumors with *MTAP* deletion, which co-occurs with *CDKN2A/B* loss, leading to the development of MTA-cooperative PRMT5 inhibitors. To identify additional collateral damage context/target pairs for chromosome 9p and other common loci of chromosomal deletions, we conducted a combinatorial CRISPR screen knocking out frequently co-deleted genes in combination with a focused target library. We identified the gene encoding the ribosome rescue factor PELO as synthetic lethal with loss of gene encoding the SKI complex interacting protein FOCAD, which is frequently co-deleted alongside *MTAP* and *CDKN2A/B* on chromosome 9p. A genome-wide screen in *FOCAD* isogenic cells further identified the ribosome rescue GTPase and PELO binding partner HBS1L as the top synthetic lethal target for *FOCAD* loss. Analysis of publicly available data and genetic manipulation of *HBS1L* using orthogonal modalities validated this interaction. HBS1L dependency in *FOCAD*-deleted cells was rescued by *FOCAD* re-expression, and *FOCAD* intact cells could be rendered HBS1L-dependent by *FOCAD* knockout, demonstrating the context specificity of this interaction. Mechanistically, *HBS1L* loss led to translational arrest and activated the unfolded protein response in *FOCAD*-deleted cells. *In vivo*, *HBS1L* deletion eliminated growth of *FOCAD*-deleted tumors. Here we propose a model where the FOCAD/SKI complex and HBS1L/PELO work together to resolve aberrant mRNA-induced ribosomal stalling, making the HBS1L/PELO complex an intriguing novel target for treating *FOCAD*-deleted tumors.

## INTRODUCTION

Deletion of tumor suppressor genes in cancer often results in co-deletion of nearby genes as collateral damage. Loss of the tumor suppressor gene itself or neighboring collateral damage genes can lead to reprogramming of pathways within the cancer cells, creating novel cancer-specific dependencies for therapeutic intervention. One example of a collateral damage context leading to synthetic lethality is loss of the *MTAP* gene, which is frequently co-deleted with the tumor suppressor genes *CDKN2A/B* due to its proximity on chromosome 9p21, causing hyperdependence on the arginine methyltransferase PRMT5 for cell survival.^1–4^ The synthetic lethal relationship between *MTAP* and *PRMT5* has served as the foundation for the translation of PRMT5 inhibitors into the clinic, where patients are currently benefiting from their use.^5–11^

The chromosome 9p21 locus harbors deletions in ∼15% of human cancer cases.^12^ Many tumor types that commonly have deletions in this region are areas of high unmet medical need, including pancreatic cancer, glioma, and non-small cell lung cancer. These cancers require novel therapeutic strategies to improve patient outcomes. We therefore sought to identify additional synthetic lethal targets focusing on chromosome 9p21 loss and other loci of common chromosomal deletions. Through a series of unbiased genetic screens, we identified a strong interaction between *FOCAD*, a gene frequently deleted as collateral damage to *CDKN2A/B* deletion, and two genes encoding proteins that work as a requisite binding pair: *HBS1L* and *PELO*. The protein product of the *FOCAD* gene interacts with the SKI (Superkiller) complex and exosome to degrade aberrant mRNAs and maintain mRNA homeostasis.^13,14^ *HBS1L* and *PELO* encode proteins that work together to rescue ribosomes that become stalled on aberrant mRNAs.^15,16^ This finding is consistent with recent work elucidating SKI complex-based synthetic lethal relationships.^17,18^

Here, we present identification and validation of the synthetic lethal interaction between *FOCAD* and *HBS1L* at the DNA and protein level. We show that the viability of *FOCAD* deleted cells depends on HBS1L, and that loss of *HBS1L* leads to destabilization of PELO protein. Inactivation of *HBS1L* leads to a decrease in translation and activation of the unfolded protein response, ultimately leading to cell death in *FOCAD*-deleted, but not *FOCAD* intact, cells. Together, these data support a model in which loss of *FOCAD* leads to buildup of aberrant mRNAs on which ribosomes become stalled^15^, and results in hyperdependence on HBS1L/PELO for ribosome rescue. Targeting the HBS1L/PELO protein complex is thus an attractive novel strategy for treating *FOCAD* deleted tumors.

## MATERIALS AND METHODS

Information for compounds, antibodies, and cell lines used in this study are listed in Supplementary Table S1.

### Cloning of sgRNA and cDNA constructs

sgRNA and cDNA constructs are based on the 3^rd^ generation lentiviral system. Sequences for sgRNAs and cDNAs are reported in Supplementary Table S4. sgRNAs and cDNA inserts were synthesized by Integrated DNA Technologies (Newark, NJ). For sgRNA cloning, annealed oligonucleotides were cloned into the appropriate destination vector using T4 DNA ligase (New England Biolabs, M0202L). For cDNA constructs, cDNA inserts were cloned into destination vectors by restriction enzyme digestion and ligation. Sanger sequencing was used to verify the fidelity of all plasmid constructs.

### Cell culture and cell line engineering

All cell lines were cultured and maintained in a humidified environment at 37°C under 5% CO_2_. Cell lines were tested for mycoplasma weekly using the Lonza MycoAlert Detection Kit (Lonza, LT07-318). Cell line information is listed in Supplementary Table S2. Cell lines were authenticated by short tandem repeat at LabCorp (Burlington, NC). Cells were used at or below passage 15 and within 8 weeks of thawing.

Lentivirus was produced by transfecting plasmid into Lenti-X cells (Takara Bio, 632180). Transfection components included the plasmid, lentiviral packaging mix (Cellecta, CPCP-K2A), Lipofectamine 3000 (Thermo Fisher Scientific, L30000015), and Opti-MEM (Gibco, 31985-062). Media was replaced with DMEM + 30% FBS after 16-24 hours. Media was collected 48 hours post-transfection and filtered through a 0.45 µm membrane.

For infection, media containing 8 µg/mL polybrene (Sigma-Aldrich, TR-1003-G) was added to seeded cells along with virus. After 16-24 hours the media was refreshed. Selection began 48 hours after infection with the antibiotic that corresponded to the vector (blasticidin (Gibco, A1113903), puromycin (Gibco, A1113803), or geneticin (Gibco, 10-131-035)).

For the generation of dTAG single-cell clones, 1000-2000 cells were seeded in 15 cm tissue culture treated dishes. Cells were cultured until singular colonies were visible. Cells were washed with PBS and trypsin-soaked filter discs were used to detach colonies. Discs were placed into individual wells of a 24-well plate containing cell culture media. For all other single cell clones, cell line pools were diluted using limiting dilution and seeded into 96-well plates at ∼1 cell/well. Colonies were expanded and genomic DNA was extracted using the DNeasy Blood and Tissue Kit (Qiagen, 69506) or the QuickExtract DNA kit (Lucigen, QE09050), according to manufacturer’s instructions. PCR was performed around the sgRNA target site using Q5® Hot Start High-Fidelity protocol (New England Biolabs, M0491). PCR products were sequenced at GENEWIZ (Boston, MA).

### CRISPR screening

CRISPR screening was conducted as reported previously^19^ with some modifications. Single cell clones were generated for both enCas12a and SpCas9 screens.

For the simultaneous and combinatorial knockout of co-deleted context genes and target genes, we engineered the enCas12a vector to include an array of 3 sgRNAs after 3 direct repeat variants as previous reported.^20^ In the 1^st^ position of the sgRNA array, one pre-validated sgRNA targeting each of 20 frequently co-deleted genes and 2 intron-cutting sgRNA controls were added individually. For the 2^nd^ and 3^rd^ positions, pairs of sgRNAs targeting the same gene were designed based on previously published designs to maximize editing efficiency^20^. For each of the 5137 candidate targets selected from a ‘druggable genome’ library, either 2 or 4 pairs of gRNAs were included for each gene in addition to intron cutting and negative control (non-cutting) sgRNAs (gene list and sgRNA sequences available in **Supplementary Dataset 1**). For the SpCas9 whole genome library, sgRNA design was based on the Brunello library^21^, with additional intron cutting and negative control (non-cutting) sgRNAs added (gene list and sgRNA sequences available in **Supplementary Dataset 2**).

The library oligonucleotide pools were synthesized at Twist Biosciences (San Francisco, CA). The synthesized products were PCR-amplified before purification and cloning into the destination vector using Golden Gate Assembly with Esp3I (NEB, R0734L) per manufacturer’s instruction. The ligated products were then column purified (Zymo Research, D4014) and electroporated into Endura ElectroCompetent cells (Bioresearch Technologies, 60242-2) per manufacturer’s instructions. The transformed cells were plated on 245 mm^2^ bioassay agar dishes (Teknova, L6010) and grown at 32 °C overnight. Colonies were maxi-prepped to obtain the final libraries.

The plasmid pool was transfected as described above using 2- or 5-layer CellSTACKs (Corning, 3313). Virus was aliquoted for storage at −80°C until use.

For library screening, target enCas12a or SpCas9 clones were infected with the respective library to obtain a multiplicity of infection of ∼40% and were selected with puromycin. Cells were passaged routinely for 14 days post-selection. gDNA was extracted from the cell pellets using the HSM 2.0 Instrument (Promega, A2715) and subjected to parallel PCR using KAPA HiFi HotStart ReadyMix (Roche, KK2602,) to amplify the regions surrounding the gRNAs in the library vectors. The PCR products were purified using KAPA Pure Beads (Roche, KK8002) and loaded onto Illumina Nexseq 2000 system (Illumina, San Diego, CA) for sequencing. Combinatorial CRISPR gene interaction analysis was performed based on the Bliss independence model^22^ (**Supplementary Fig. S2**), in which the expected effect of sgRNA pair was predicted by an additive quadratic fit of the effects of the single sgRNAs composing the sgRNA pair^23^. Observed effect of each sgRNA pair was measured by the log2 fold change (log2FC) of its read count at end of experiment compared to its read count in plasmids. Effect of Genetic interaction (GI) was calculated by subtracting the expected effect from the observed effect for each sgRNA pair^23^. Raw GIs were then corrected by dispersion adjustment^24^, normalized by the standard deviation of the 1000 nearest neighbors in terms of the expected effect^25^ . The adjusted GI values were further converted to z-scores of sgRNA pairs (zGI) and two-tailed p-values were derived by z-test standardization. Finally, the average of zGIs across all sgRNA pairs targeting co-deleted gene A and druggable library gene B was defined as the gene pair score (GPS_AB_) and the p-value of GPS_AB_ was derived using Stouffer’s method^26^. Threshold levels of significance were adjusted for multiple tests by Bonferroni correction (0.05/(5137*20) = 5×10^−7^). Details of the combinatorial CRISPR gene interaction analysis were described in Supplementary Methods.

### DepMap FOCAD expression vs target gene CRISPR score analysis

CRISPR dependency scores estimated by Chronos and processed RNAseq data of CCLE cell lines were downloaded from DepMap (2022q2)^27^ . We tested the correlation between each target gene and FOCAD expression by linear regression. Cell line lineage was adjusted in each linear test, the RNA-seq gene expression value (log2TPM+1) of FOCAD gene was used as the predictor variable and normalized CRISPR scores of gene dependency from DepMap were used as the response variable to fit a linear regression model.

### Colony formation assay

Cells were seeded into tissue culture treated 6-well plates and allowed to adhere overnight (MIAPACA2 at 1,000-1,200 cells/well; T24 at 400 cells/well; CFPAC1, LN229, and LN18 at 2,000 cells/well; ACHN at 3,000 cells/well) . For dTAGv1 treatment, each well was dosed with dTAGv1 (R&D Systems, 6914) with DMSO vehicle for control. After initial dosing, dTAGv1-containing media was replaced every 2 days for 14 days. For doxycycline treatment, cell culture media +/-0.5 mg/mL doxycycline (R&D Systems, 4090) was refreshed every 2-3 days. For experiments in HeLa wild-type and *FOCAD* knockout cell lines, cells were seeded at 5,000 cells per well into 6-well plates (ThermoFisher, 140675) and allowed to adhere overnight. Cells were then infected with the sgITC or sgHBS1L lentivirus. Blasticidin (ThermoFisher, A1113903) selection at 8 µg/ml was initiated 48 hours post-infection. For all colony formation experiments, the cells were washed with PBS at endpoint and stained with 1% crystal violet solution (Sigma-Aldrich, V5265). After 5 minutes rocking, the crystal violet was removed and cells were washed by submerging the plate in water. The plates were dried overnight and imaged using either the LI-COR Odyssey Clx and Image Studio software (LI-COR) or Canon photo scanner. For quantification, 500 μL 10% acetic acid solution was added and rocked 30 minutes. Absorbance was measured in a 100 μL aliquot at 570 nm on the SpectraMax M5 (Molecular Devices).

### Immunoblotting

Protein lysate was prepared by scraping cells in RIPA buffer (Life Technologies, 89901) supplemented with protease and phosphatase inhibitors (Life Technologies, 78445) and universal nuclease (Life Technologies, 88700) or EBC buffer (Boston Bioproducts, BP-1410) supplemented with Halt Protease Inhibitor (ThermoFisher, 78425). Lysates were clarified by centrifugation at maximum speed for 10 minutes at 4°C and protein concentration was determined (Pierce 660 nm Protein Assay, Thermo Scientific, 22660).

For traditional western blotting, 20-40 μg of protein was denatured in LDS sample buffer (Invitrogen, NP0007) containing DTT at 95°C for 5 minutes. DNase digestion was performed on all samples by adding 1 μL DNase I (Invitrogen, 18047019) and incubating at 37°C for 30-60 minutes. Samples were separated by SDS-PAGE electrophoresis in 4-12% Bis-Tris gels (Thermo Scientific, WG1402BOX). Protein was transferred to nitrocellulose membranes (Thermo Scientific, STM2008) using NuPAGE Transfer Buffer (Thermo Scientific, NP00061) supplemented with NuPAGE Antioxidant (Thermo Scientific, NP0005) and 20% methanol.

For simple western, 1.2-1.8 μg protein was denatured in 0.1X sample buffer (ProteinSimple, 042195) supplemented with DTT and fluorescent master mix for Jess (ProteinSimple, PS-FL01-8). Samples were loaded onto Jess plates and processed according to the manufacturer’s protocol.

### Puromycin incorporation assay

To determine translational arrest, cells were plated at a density to achieve 70-80% confluency at endpoint. Cells were treated with indicated compounds for indicated timepoints. Prior to puromycin labeling, media was removed and cells were washed with 1X PBS (Invitrogen, 10010001). Puromycin (20µM) (Gibco, A1113803) was mixed with pre-warmed media, added directly to cells, and incubated at 37 °C for 15 minutes. Cells were washed with ice-cold 1X PBS and directly lysed in plate using ice-cold RIPA buffer (Thermo Fisher, 89900) supplemented with 1X Halt Protease and Phosphatase Inhibitor (Thermo Fisher, 78446). Following protein extraction, immunoblotting was performed where equal amounts of protein lysates were loaded on 4-12% Bis-Tris protein gels (Thermo Fisher, WG1403) and puromycin incorporated nascent peptides was identified using an anti-puromycin antibody (Sigma Aldrich, MABE343). The intensity of Puromycin was normalized to ponceau (Thermo Fisher, A40000279) loading control.

### qPCR

Cells were plated to achieve 80% confluency at collection and allowed to attach overnight. Cells were treated with DMSO, 10 nM, 100 nM or 1 μM dTAGv1 (R&D Systems, 6914) and incubated for 24 hours. After incubation, cells were washed once with DPBS, then lysed in RNA Cell Lysis Buffer (Boston Bioproducts, R-108) supplemented with 1:40 RNasin Recombinant Ribonuclease Inhibitors (Promega, N2515). mRNA levels were detected using Taqman Fast Virus 1-Step Master Mix (Life Technologies, 4444436) according to manufacturer’s protocol.

### Cell cycle assay

Cells were seeded on 6-well plates (Invitrogen, 140675) and compound was administered for the specified time point and concentration. Cells were then harvested, resuspended in PBS (Invitrogen, 10010001), pelleted, and washed twice to eliminate cellular debris. Cells were fixed in 70% ethanol (Invitrogen, T038181000), stained with a FxCycle PI/RNase Staining Solution (20 µg/mL PI and RNaseA diluted in PBS) (Invitrogen, F10797) and subjected to flow cytometry analysis using a Novocyte Flow Cytometer (Agilent, Santa Clara, CA). At least 20,000 live cells were counted for each biological replicate. Flow cytometry data was analyzed using FlowJo (FlowJo, LLC, Ashland, Oregon). Live cells were gated to remove any sub-G1 cells and then were subjected to cell cycle analysis via DeanJett-Fox modeling.

### Xenograft model in NOG mice

*In vivo* studies were conducted at Pharmaron (China) in accordance with AAALAC guidelines and using an IACUC-approved protocol. Female 6-8 week old NOG mice (Vital River Biotech Co., Ltd.) were subcutaneously inoculated on one flank with 5×10^6^ MIAPACA2 cells engineered to express Cas9, a doxycycline-inducible cDNA encoding the *HBS1L* gene, and either an intron-targeting or *HBS1L*-targeting sgRNA in a 1:1 suspension with Matrigel and 0.5 μg/mL doxycycline. Mice were given free access to Dyets doxycycline chow (650 ppm doxycycline) until tumor volumes reached an average of 150 mm^3^. Mice were then randomized to either continue receiving Dyets doxycycline chow (+DOX arms) or to be switched to Bio-Serv normal chow (vehicle arms). Tumor measurements were conducted using calipers twice per week and animals were sacrificed after the indicated number of days of treatment and their tissues harvested for assessment of pharmacodynamic markers.

### Statistical analysis

ImageJ was used to quantify images (National Institutes of Health, Bethesda, MD). GraphPad Prism (GraphPad, Software Inc., La Jolla, CA) was used for all statistical analyses.

### Data availability statement

Data described in this manuscript are presented within the article and the Supplementary Materials. Additional information can be requested by contacting the corresponding author.

## RESULTS

### Combinatorial CRISPR screens identified multiple mRNA homeostasis factors as synthetic lethal targets in *FOCAD*-deleted cells

To identify novel synthetic lethal targets associated with collateral damage arising from tumor suppressor gene loss, we took advantage of the enhanced Cas12a (enCas12a) system,^20,28^ which allows for simultaneous and high throughput expression of multiple sgRNAs from a single vector enabling the combinatorial knockout of gene pairs. We interrogated a library of >5000 candidate target genes (enriched for genes encoding druggable targets) against 20 context genes frequently found to be co-deleted with tumor suppressor genes (**Fig.1A, B**). In the first position of the enCas12a gRNA plasmid we cloned a pre-validated gRNA targeting one of the frequently co-deleted genes (**Supplementary Fig. S1)**. In the second and third positions of the plasmid we installed pairs of gRNAs targeting the same gene to enhance editing efficiency (**Fig.1B**). Overall, we systematically examined >100,000 genetic interactions across 4 cell lines representing different lineages. With built in intron-targeting control gRNAs, we were able to examine the single gene knockout effect of both the co-deleted context genes and each of the >5,000 candidate target genes. We then used gene interaction analysis to calculate the difference between the observed effect of combined knockout versus the estimated additive effect of the individual gene knockouts. Then, we quantified any synergistic interaction between the context and target genes using the Bliss independence model as measured by log2 fold change (log2FC). These values were converted to z-scores, and significance was calculated using Stouffer’s method (full method and calculations available in Methods Section and Supplementary Materials **Supplementary Fig. S2**).

**Figure 1.**
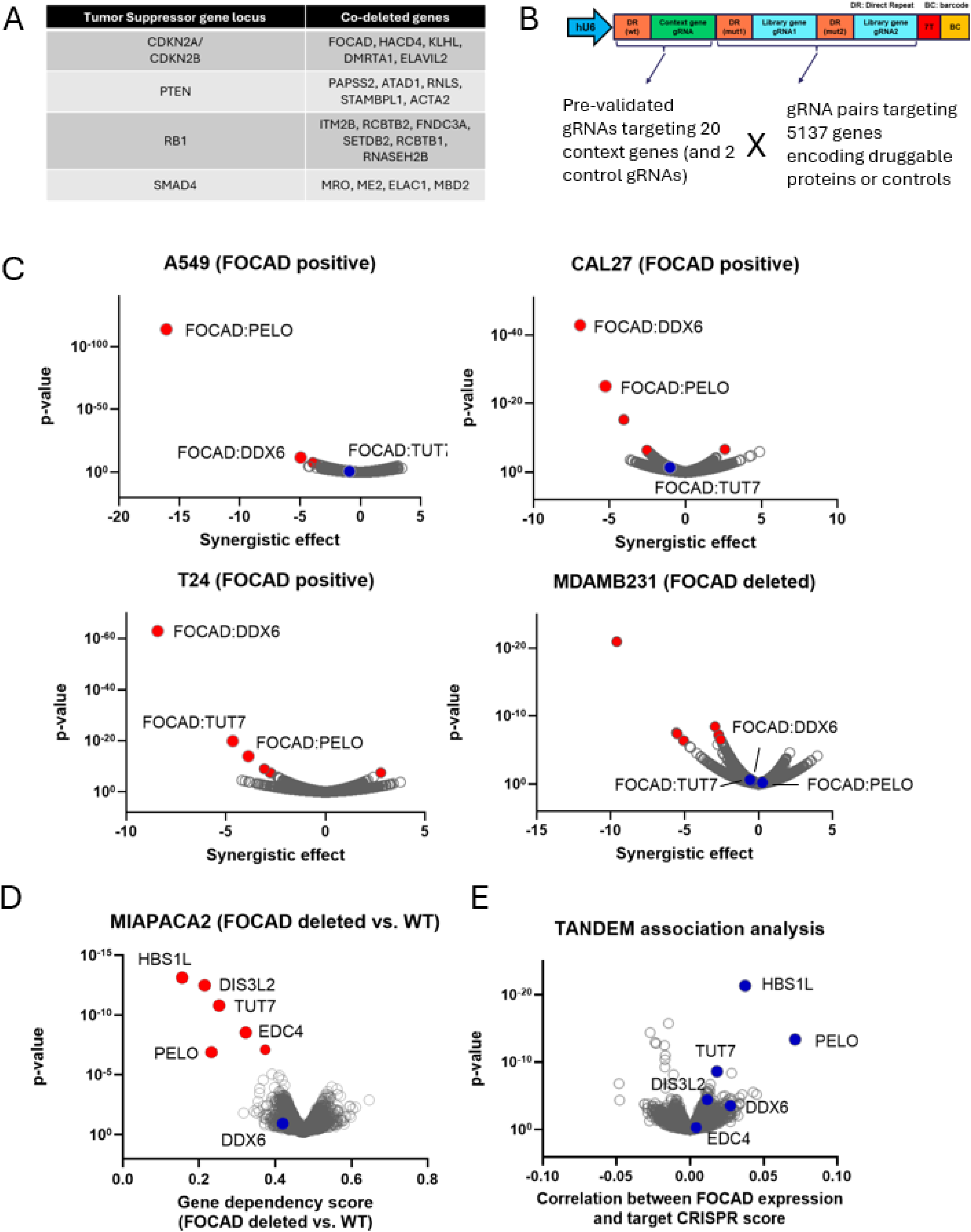
Orthogonal CRISPR screens identified *HBS1L/PELO* as the top synthetic lethal targets for *FOCAD*-deleted context. A,B) enCas12a combinatorial CRISPR library design (B) for investigating synthetic lethal targets for 20 co-deleted genes surrounding frequently deleted tumor suppressor genes (A). C) Volcano plots of the enCas12a combinatorial screen results examining ∼100K genetic interactions in the indicated cell lines. X-axis: calculated synergistic score where negative values indicate synergy between two evaluated genes. Y-axis: *p*-value, calculated using Stoufer’s method combining guide level p-values of quadratic regression model of gene-gene interactions. Significant gene pairs are highlighted as red dots. D) Volcano plots of the genome-wide Cas9 screen in engineered *FOCAD* isogenic MIAPACA2 cell lines. X-axis: calculated gene dependency score. Value<0.5 indicates greater target gene dependency in *FOCAD*-deleted cell line versus *FOCAD* intact cell line. Y-axis: *p*-value, calculated using Unique Molecular Identified Bayesian Beta-binomial (UMIBB) test. Significant gene hits are highlighted as red dots. E) Volcano plots of the TANDEM analysis for association between target gene CRISPR score and *FOCAD* gene expression across the Achilles screen cell line panel^31^ . Value>0 indicates positive correlation and potential targets for *FOCAD*-deleted tumors. *p*-values were calculated using linear regression tests.

Among all the combinations tested, multiple mRNA surveillance and homeostasis factors, including *PELO* and *DDX6*, consistently emerged as top synthetic lethal hits in the *FOCAD*-deleted context across 3 cell lines that endogenously express *FOCAD* (A549, CAL27, T24), but not the endogenously *FOCAD-*deleted cell line MDA-MB-231 (**Fig. 1C**, full results in **Supplementary Dataset 3**). In contrast, the screen in the MDA-MB-231 cell line, which already has *FOCAD* inactivated, showed no additional effect of expressing the *FOCAD* sgRNA and thus serves as an independent cell line control for synergy calculations. In models that endogenously express *FOCAD*, knockout of *FOCAD* alone has little effect on viability, but combined knockout of *FOCAD* and either *PELO* or *DDX6* exceeds the expected additive effect indicating a synergistic interaction between the genes (**Supplementary Fig. S2**). In T24 cells we also identified the Terminal Uridylyl Transferase 7 (*TUT7*) and *FOCAD* as a top scoring genetic interaction, consistent with recent studies evaluating the synthetic lethality between these two genes and the role of TUT7 in regulating RNA stability through the addition of uridine residues (U-tails) to the 3’ end of RNA molecules.^29^ Given the prevalence of *FOCAD*-based gene pairs identified at hits in this initial screen, we decided to conduct a genome-wide CRISPR screen to identify the full set of *FOCAD* synthetic lethal interactions.

### Genome wide Cas9 screen in *FOCAD* isogenic cell line pair identified PELO binding partner HBS1L as the top synthetic lethal target for *FOCAD*-deleted cells

To confirm the combinatorial screen results with an orthogonal unbiased screen and expand the target space, we carried out genome-wide SpCas9 CRISPR screens in a pair of engineered isogenic cell lines that differ only by *FOCAD* expression through stable infection with either *FOCAD* cDNA or an empty control vector in the *FOCAD*-deleted MIAPACA2 cell line (**Fig. 1D**, **Supplementary Fig. S3**, full results in **Supplementary Dataset 4**). UMIBB analysis^30^ of the results confirmed *PELO* and *TUT7* among the top synthetic lethal hits with *FOCAD* loss. More interestingly, *HBS1L*, which encodes a ribosome rescue GTPase and uses PELO as a requisite binding partner, was identified as the top synthetic lethal interaction with *FOCAD* loss (**Fig. 1D**). This interaction was stronger than that of *PELO*:*FOCAD*, but was not identified in the initial combinatorial screen as *HBS1L* was not included in the sgRNA library. To further support our screen results and provide direction for prioritization, we leveraged the internal Tango dependency map (TANDEM) analysis on the Achilles screen dataset^31^, in which we calculated the association of the CRISPR score of each target gene with the mRNA expression of *FOCAD* and generated waterfall plots assessing bimodality of target gene dependence based on *FOCAD* status as well as essentiality (**Fig. 1E**, **Supplementary Fig. S4**). *HBS1L* CRISPR score showed the best correlation with *FOCAD* expression and copy number out of all genes, consistent with our unbiased screening approach. Further, *PELO* showed strongly negative CRISPR scores across the majority of cell lines, suggesting it may be an essential gene and therefore have a limited therapeutic index. For these reasons, we prioritized *HBS1L* for further validation.

### Multiple orthogonal genetic manipulation approaches validate *HBS1L* as a synthetic lethal dependency for cell lines with *FOCAD* loss *in vitro*

We next carried out single gene *in vitro* validation studies to ask whether the synthetic lethal relationship between *HBS1L* and *FOCAD* was reproducible outside the screen setting. To determine the impact of complete *HBS1L* loss we first expressed a doxycycline-inducible CRISPR-resistant *HBS1L* cDNA in Cas9-expressing MIAPACA2 cells, which endogenously lack *FOCAD*. While keeping cells cultured with doxycycline to maintain exogenous *HBS1L* expression, we introduced either an intron cutting control (ITC) sgRNA or two different sgRNAs targeting endogenous *HBS1L* to create models in which *HBS1L* expression is controlled by doxycycline (**Fig. 2A**). Immunoblot analysis confirmed the loss of HBS1L upon doxycycline withdrawal in the models with endogenous *HBS1L* knockout (**Fig. 2B**). Loss of HBS1L also resulted in depletion of PELO, possibly due to lack of stability of PELO in the absence of HBS1L in keeping with previous studies.^32–34^ Consistent with our screen results, colony formation was impaired when exogenous *HBS1L* expression was removed (doxycycline withdrawn) in MIAPACA2 models lacking endogenous *HBS1L* (**Fig. 2C**). There was no significant impact of removing exogenous *HBS1L* expression in cells with intact endogenous *HBS1L*. These data demonstrate that *FOCAD*-null MIAPACA2 cells are hyperdependent on HBS1L for survival, and that this effect is on-target as it could be rescued by exogenous expression of *HBS1L*.

**Figure 2.**
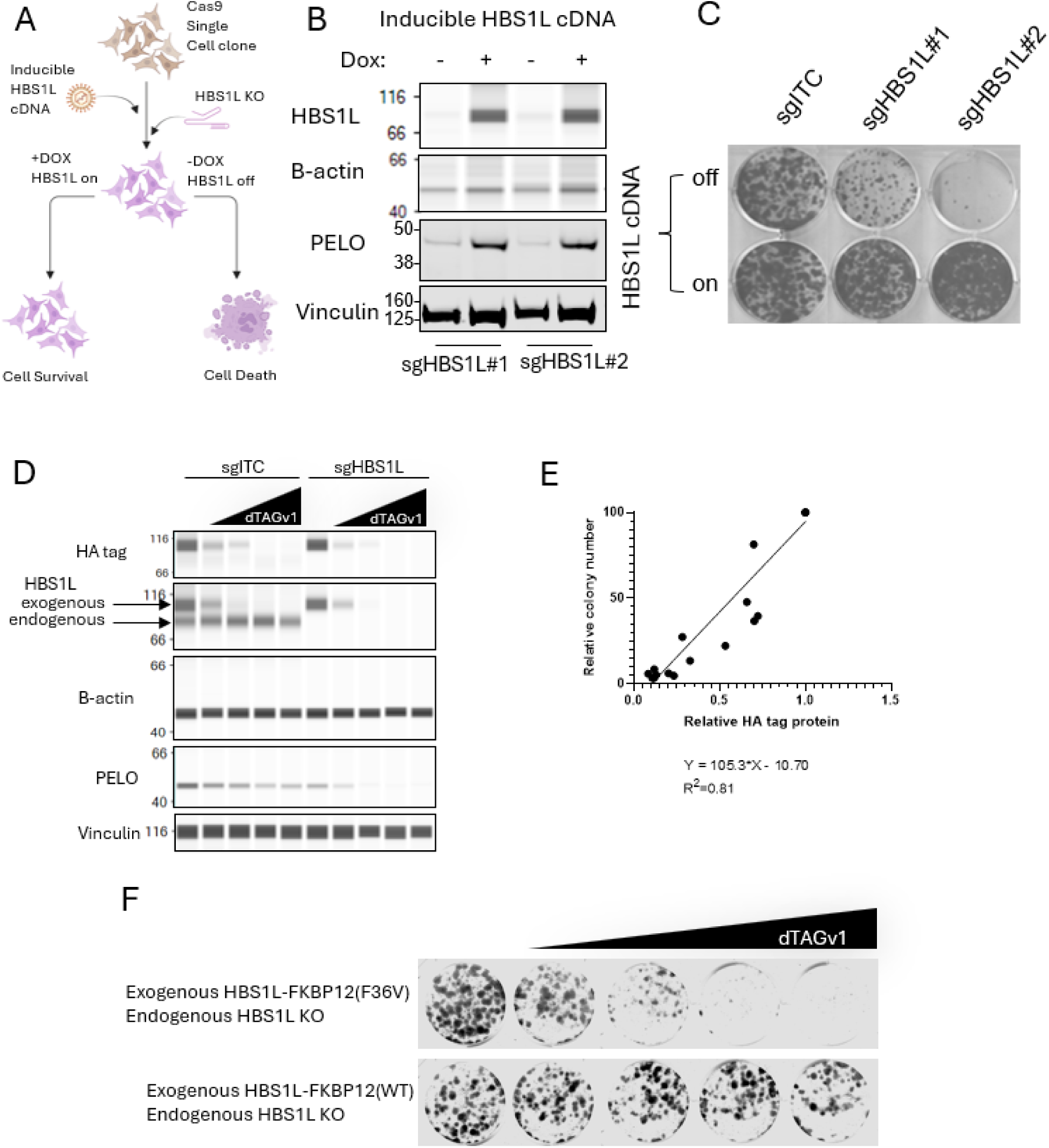
HBS1L is a synthetic lethal target in *FOCAD*-deleted MIAPACA2 cells. A) Scheme of *HBS1L* genetic validation scheme with exogenous *HBS1L* cDNA and endogenous *HBS1L* knockout. B) Western blot of pooled MIAPACA2 cells stably expressing doxycycline inducible CRISPR resistant *HBS1L* cDNA and constitutive *HBS1L* sgRNAs. C) Colony formation assay of pooled MIAPACA2 cells stably expressing doxycycline inducible CRISPR resistant *HBS1L* cDNA and either intron cutting control sgRNA or the constitutive *HBS1L* sgRNAs. Data are representative of N=3 biological replicates. D) Western blot of a clonal population of MIAPACA2 cells stably expressing *HBS1L-FKB12(F36V)* fusion cDNA and having endogenous knockout of *HBS1L* treated with 0, 1, 5, 20, or 50 nM dTAGv1 for 4 days. E) Correlation of relative colony number of the MIAPACA2 clone stably expressing *HBS1L-FKB12(F36V)* fusion cDNA and endogenous *HBS1L* knockout in the long-term colony formation assay in F) versus relative HA-tagged HBS1L expression as measured in western blot in D). F) Colony formation assay of clonal populations of MIAPACA2 cells stably expressing either *HBS1L-FKBP12(G12V)* or *HBS1L-FKBP12WT* fusion cDNA and having endogenous knockout of *HBS1L* treated with 0, 0.5, 1, 5, 20, or 50 nM dTAGv1 for 14 days. Data are representative of 7 single cell clones.

To ask whether the results of manipulating *HBS1L* at the DNA level would be reproducible when manipulating HBS1L at the protein level, we engineered MIAPACA2 cells to express an edit-resistant *HBS1L* cDNA fused to an HA tag and either the FKBP12(F36V) degron (HBS1L-dTAG) or a non-degradable control degron (FKBP12 WT)^35,36^ covering endogenous *HBS1L* knockout (or control cells in which an intron cutting control sgRNA (sgITC) was used in place of the *HBS1L* targeting sgRNA (sgHBS1L)). We then generated single cell clones and confirmed complete knockout of endogenous *HBS1L* in the sgHBS1L clones (7 independent clones). Administering different dosages of dTAGv1 ligand allowed titration of HBS1L protein levels in the HBS1L-FKBP12(F36V) cells (**Fig. 2D, E**), but did not affect the HBS1L-FKBP12 WT fusion protein (**Supplementary Fig. S5A**). Depletion of HBS1L protein resulted in a dose-dependent decrease in colony formation, further highlighting the dependence of *FOCAD*-deleted MIAPACA2 cells on HBS1L for survival (**Fig. 2F**). Consistent with our DNA level manipulation, loss of HBS1L at the protein level also led to a decrease in PELO expression (**Fig. 2D**).

To expand upon our results in MIAPACA2 cells, we generated a small panel of three cell lines having intact *FOCAD* (T24, CFPAC1, LN229) and three cell lines with endogenous loss of *FOCAD* (ACHN, MIAPACA2, LN18) and asked whether each was dependent on *HBS1L* for survival. We expressed a CRISPR edit-resistant doxycycline-inducible *HBS1L* cDNA and knocked out endogenous *HBS1L* in each model, then performed colony formation assays in the presence (on) or absence (off) of HBS1L (with or without doxycycline, respectively) (**Fig. 3A**). All three cell lines lacking functional *FOCAD* failed to produce robust colonies upon *HBS1L* loss, while all three cell lines with intact *FOCAD* showed similar growth in the presence or absence of *HBS1L*. HBS1L depletion was confirmed at the protein level (**Supplementary Fig. S5B**).

**Figure 3.**
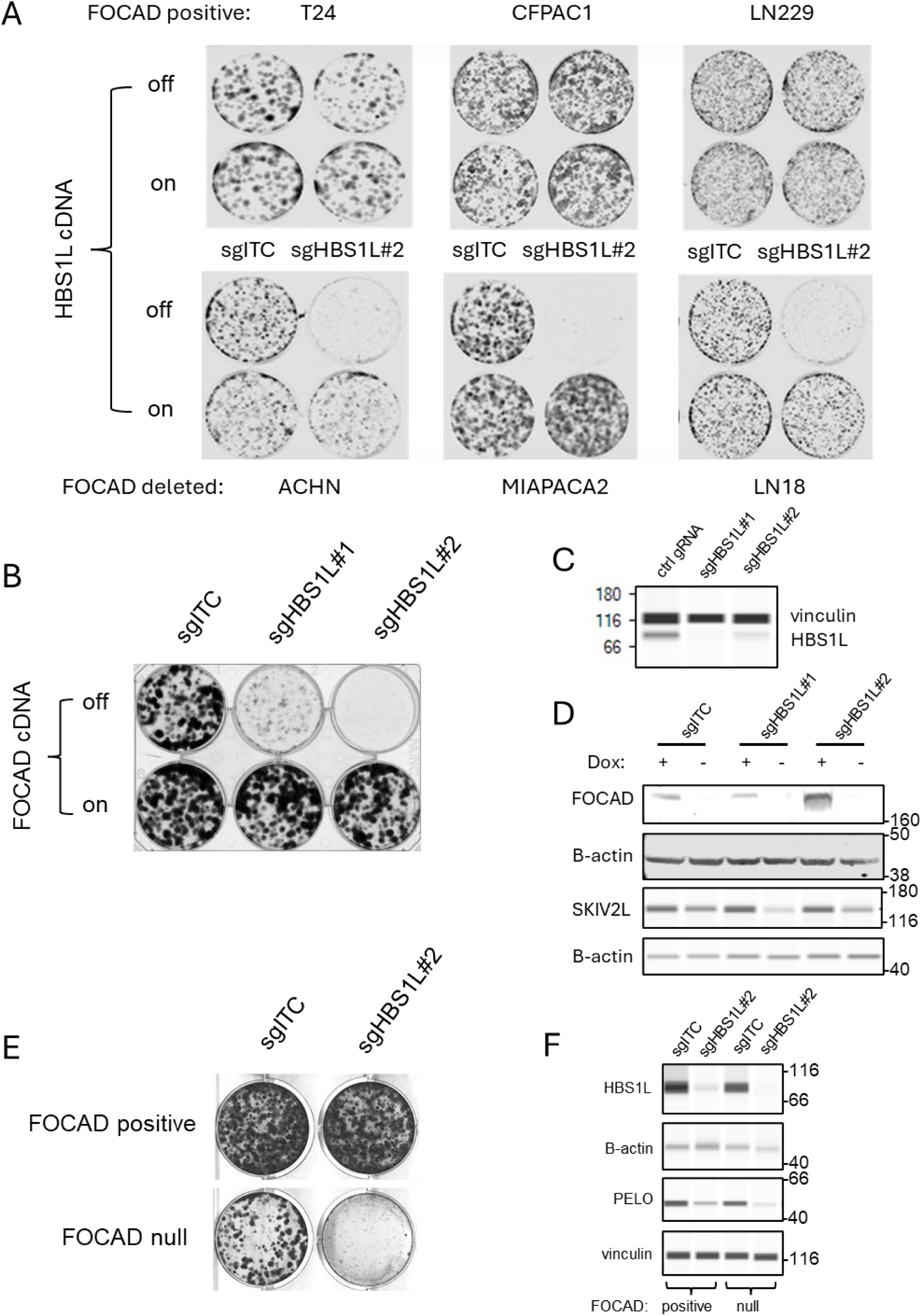
HBS1L dependency relies on FOCAD status. A) Colony formation assays in a panel of cell lines with indicated FOCAD status stably expressing a doxycycline inducible CRISPR resistant *HBS1L* cDNA and harboring endogenous HBS1L knockout. Data are representative of N=2-3 biological replicates. B) Colony formation assay of MIAPACA2 cells expressing a doxycycline inducible *FOCAD* cDNA and harboring endogenous HBS1L knockout. Data are representative of N=3 biological replicates. C) Western blot of HBS1L in cell models shown in B) cultured in 0.5 mg/mL doxycycline. D) Western blot of FOCAD and SKIV2L in cell models shown in B) cultured with and without 0.5 mg/mL doxycycline. E) Colony formation assay of HeLa parental or *FOCAD* knockout cells expressing either an intron cutting control sgRNA (sgITC) or a sgRNA targeting HBS1L (sgHBS1L#2). Data are representative of N=3 biological replicates. F) Western blot of HBS1L and PELO in models shown in E). All western blot data in Figure 3 are representative of N=2 biological replicates.

To conclude our *in vitro* validation efforts, we expressed a doxycycline-inducible *FOCAD* cDNA into the endogenously *FOCAD*-deleted MIAPACA2 cell line and then either did (sgHBS1L) or did not (sgITC) knock out *HBS1L* (**Fig. 3B-D**). FOCAD expression was confirmed by western blot (**Fig. 3D**) and its stabilization of the SKI complex was confirmed by corresponding expression changes in SKI complex protein SKIV2L. Growth of *HBS1L-*deficient MIAPACA2 cells was completely abrogated without *FOCAD* expression (off), and was completely restored upon expression of the *FOCAD* cDNA (on). Conversely, we examined the impact of *FOCAD* loss on HBS1L dependency in a cell line that is endogenously *FOCAD* proficient and carried out similar assay. To achieve this, we took advantage of the commercially available isogenic matched wild-type and FOCAD knockout HeLa cells from Abcam. As expected, the loss of *FOCAD* rendered HeLa cells dependent on HBS1L for survival (**Fig. 3E, F**, **Supplementary Fig. S5C**). Taken together, these data reveal a robust synthetic lethal interaction between *HBS1L* and *FOCAD* at both the DNA and protein level across diverse models and using multiple orthogonal methods.

### Loss of *HBS1L* eliminates growth of *FOCAD*-deficient tumors in mouse xenografts

We next sought to ask whether the synthetic lethal relationship we discovered between *HBS1L* and *FOCAD in vitro* would hold true *in vivo*. We implanted mice with xenografts of a derivative of the *FOCAD*-deleted cell line MIAPACA2 that had been engineered to express a doxycycline-inducible cDNA encoding *HBS1L* in the presence or absence of endogenous *HBS1L*. Tumors implanted in mice who were fed chow containing doxycycline, and therefore expressed exogenous *HBS1L*, grew regardless of the status of endogenous HBS1L (**Fig. 4A-D**). Similarly, tumors having intact endogenous *HBS1L* grew equally in mice who were fed chow with doxycycline (+DOX), and thus expressed both endogenous and exogenous *HBS1L*, or mice who were fed chow without doxycycline (vehicle) and thus expressed only endogenous, but not exogenous, *HBS1L* (**Fig. 4A, C-D**). In contrast, growth of tumors having endogenous *HBS1L* knocked out was abrogated in mice who were fed chow that did not contain doxycycline (vehicle), as these tumors expressed neither endogenous nor exogenous *HBS1L* (**Fig. 4B-D**). These data demonstrate that *FOCAD*-deleted tumors depend on *HBS1L* for growth, and inactivation of *HBS1L* abolishes growth of these tumors. These data were reproduced using xenografts of pooled cells having *HBS1L* knockout as well as using two independent single cell clones with confirmed complete frameshift mutations in endogenous *HBS1L* (**Fig. 1B**, **Supplementary Fig. S6**). Western blot analysis of tumors from endpoint confirmed loss of HBS1L in the relevant models, as well as destabilization of PELO upon HBS1L inactivation, consistent with our *in vitro* data (**Fig. 4C**, **Supplementary Fig. S6**).

**Figure 4.**
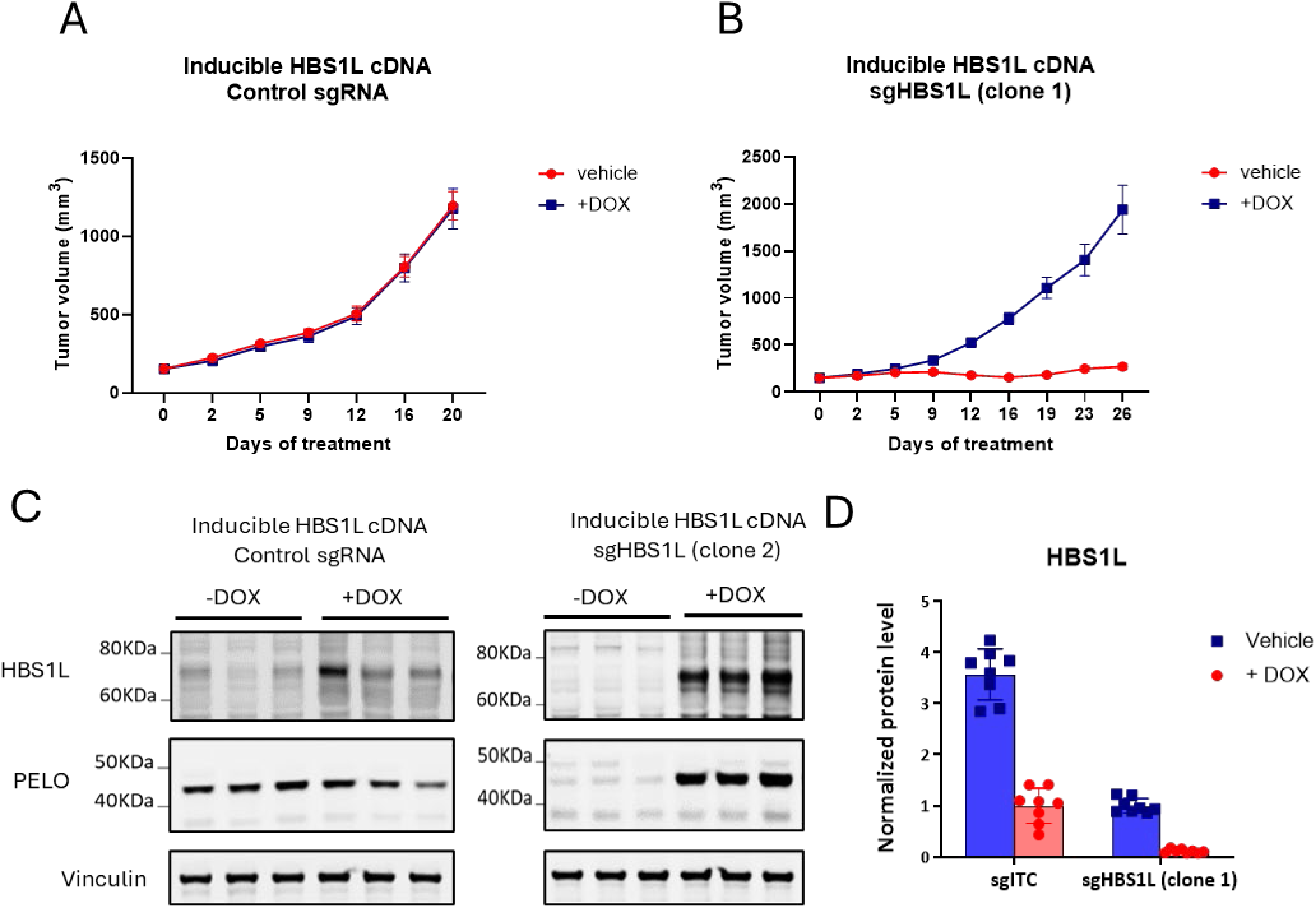
HBS1L loss eliminates tumor growth in CDX model of FOCAD-deleted pancreatic cancer. A, B) Growth of tumors in mice (N=8 animals/arm) implanted with MIAPACA2 cells expressing a doxycycline-inducible cDNA encoding HBS1L cDNA in the presence (A) or absence of endogenous HBS1L (B). In (B), implanted cells were from a single cell clone with Sanger sequencing-confirmed complete frameshift in the endogenous *HBS1L* gene. Mice were fed chow with (+DOX) or without (vehicle) doxycycline to maintain or withdraw, respectively, expression of HBS1L. Data represent the average of N=8 animals per arm, error bars represent ± SEM. C, D) Representative image of HBS1L and PELO levels (C) and quantification (D) of HBS1L level detected by Western blotting of tumors at endpoint. Data represent the average of N=8 animals per arm, error bars represent ± SD.

### *HBS1L* inactivation reduces global translation and results in cell cycle arrest and induction of the unfolded protein response

We next sought to understand the mechanistic ramifications of HBS1L inactivation in cells with *FOCAD* loss. Because of the role of HBS1L and its partner protein PELO in facilitating translation through rescue of stalled ribosomes^15,16^, we hypothesized that HBS1L inactivation, specifically in the context of FOCAD loss, would result in translational arrest. Using an *HBS1L* cDNA fused to the FKBP12(F36V) degron (dTAG)^35^, we titrated HBS1L protein level by treatment with the PROTAC dTAGv1^36^ in MIAPACA2 cells harboring endogenous *HBS1L* knockout. Using targeting protein degradation, we then asked whether reducing HBS1L level affected the ability of these cells to incorporate puromycin into growing proteins as an indicator of translation rate (**Fig. 5A-C**). Decreasing HBS1L protein level with dTAGv1 reduced puromycin incorporation in a dose-dependent manner, indicating that translation rate correlates with HBS1L level. Without proper translation, normal progression of cells through the cell cycle can become impaired. Consistently, we observed a significant increase in the population of cells arrested in G2/M phase when HBS1L was degraded using the HBS1L-dTAG model as compared to cells that retained HBS1L expression (**Supplementary Fig. 7A**), supporting our hypothesis that halted translation leads to an inability of cells to progress through the cell cycle normally.

**Figure 5.**
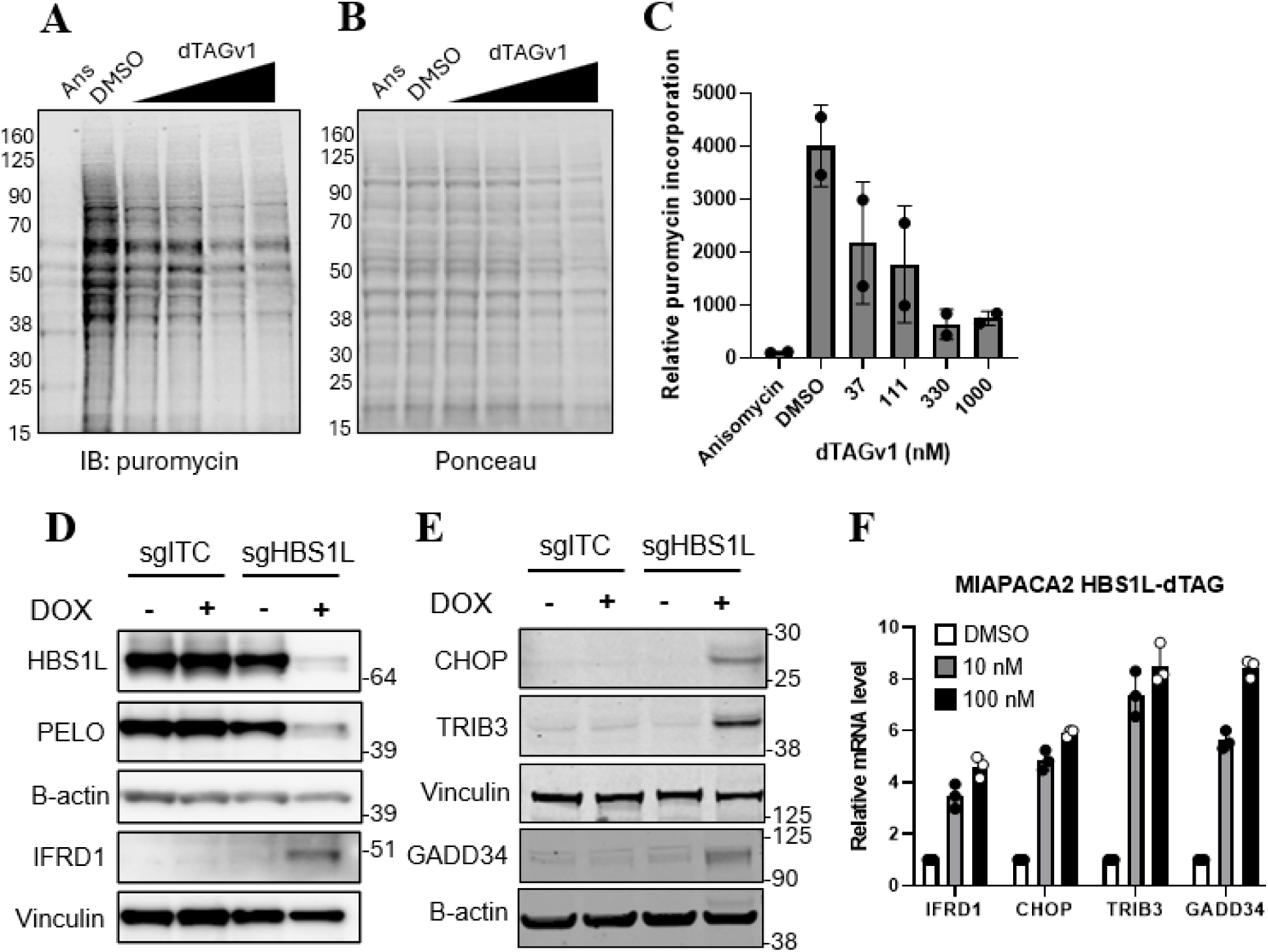
HBS1L inactivation induces translational arrest, activation of the unfolded protein response, and cell cycle arrest in *FOCAD*-deficient cells. **A-C)** Western blot of puromycin (A), Ponceau stain (B), and quantification of puromycin/Ponceau optical density (C) in MIAPACA2 cells expressing a dTAG-fused *HBS1L* cDNA covering endogenous *HBS1L* knockout and treated with the indicated doses of dTAGv1 for 5 days. Cells were pulsed with puromycin for 15 minutes prior to lysis to evaluate translational rate. Anisomycin (ANS) was used as a positive control for translation arrest. dTAGv1 dosed at 0.37, 0.11, 0.33, 1 μM. Data are representative images of 2 biological replicates (A,B) or are presented as the mean of 2 biological replicates ± SD (C). B) Ponceau stain demonstrating protein loading for western blot in (A). D,E) Western blot probing for HBS1L, PELO, and IFRD1 (D) or CHOP, TRIB3, and GADD34 (E) upon doxycycline (DOX)-inducible knockout of an intron-targeting control sgRNA (sgITC) or an sgRNA targeting HBS1L (sgHBS1L). Doxycycline was incubated for 5 days prior to sample collection. F) qPCR analysis evaluating relative level of mRNA of the indicated genes MIAPACA2 cells expressing a dTAG-fused *HBS1L* cDNA covering endogenous *HBS1L* knockout and treated with the indicated doses of dTAGv1 for 24 hours. Data represent mean of N=3 biological replicates, error bars represent ± SD.

Halted translation can result in the production of misfolded or incomplete proteins, which can activate the unfolded protein response.^37,38^ We therefore asked whether we could observe evidence of activating the unfolded protein response in response to HBS1L loss. IFRD1 is a master regulator of genes activated during endoplasmic reticulum stress, and its expression is induced upon activation of the unfolded protein response.^39^ Using a doxycycline-inducible sgRNA targeting *HBS1L* (sgHBS1L) or an intron-targeting control (sgITC), we compared IFRD1 protein level in MIAPACA2 cells that lack or have, respectively, HBS1L (**Fig. 5D**). IFRD1 expression was induced in cells in which *HBS1L* had been knocked out (sgHBS1L, +DOX), but not cells in which the sgRNA targeted an intron (sgITC). This indicates that the translational arrest observed upon HBS1L loss can activate the unfolded protein response in FOCAD-deficient MIAPACA2 cells. Consistent with these data, several genes whose increase is associated with the unfolded protein response (*IFRD1, CHOP, TRIB3, GADD34*)^40^ and their protein products were upregulated at the protein (**Fig. 5E**) and mRNA (**Fig. 5F**) levels in MIAPACA2 cells in which HBS1L had been inducibly knocked out (protein level) or degraded using an HBS1L-dTAG fusion protein (and in which endogenous *HBS1L* had been knocked out) (mRNA level). No increase in mRNA level was observed upon dTAGv1 treatment of cells expressing a non-degradable control fusion protein (HBS1L-FKBP12WT), demonstrating that the effect was not due to dTAGv1 treatment alone but rather was specific to HBS1L protein decrease (**Supplementary Fig. 7B**). Notably and consistent with our therapeutic hypothesis, using a MIAPACA2 cell line pair that was isogenic for FOCAD and expressed a DOX-inducible *HBS1L* cDNA covering endogenous *HBS1L* knockout, we observed that *CHOP* and *TRIB3* mRNA levels were induced upon HBS1L inactivation in the FOCAD-deficient MIAPACA2 cells, but not in the FOCAD-reconstituted MIAPACA2 cells (**Supplementary Fig. 7C**). These data further support the FOCAD-deletion context specificity of initiating the cellular stress response when HBS1L is inactivated.

## DISCUSSION

Precision oncology target discovery has been a recent area of focus due to the potential large therapeutic window offered by this approach, which differentiates the mutation-driven cancer cells from normal tissues. While there is a well-established clinical portfolio of drugs targeting proteins with gain-of-function mutations, exploring vulnerabilities for loss-of-function genetic contexts requires identification of novel co-dependencies such as the synthetic lethal relationship between *BRCA1* and PARP1.^41–43^ This interaction highlights how mutations in tumor suppressor genes can lead to rewiring of the cellular process, creating context-specific dependencies. As an expansion of this concept, the imprecise nature of genomic deletions (often driven by inactivation of a tumor suppressor gene at the center of the deletion locus) may result in co-deletion of neighboring non-essential genes along with the driver tumor suppressor genes. This may create unique opportunities for novel genetic dependencies, termed “collateral lethality”.^44^ The hyperdependency on PRMT5 activity in tumors having lost *MTAP*, a gene inactivated as collateral damage from the deletion of neighboring tumor suppressor genes *CDKN2A/B*, is a well-documented example of such collateral lethality.^1–4^

To identify additional targets for “collateral damage” contexts, we took advantage of the enCas12a combinatorial CRISPR platform^20,28^ to systematically survey the druggable genome space (>5,000 candidate target genes) against 20 frequently co-deleted genes surrounding the *CDKN2A/CDKN2B*, *RB1*, *PTEN* and *SMAD4* loci in four cell lines from different lineages. This platform allowed interrogation of >100,000 gene-gene interactions in a high throughput manner. Multiple factors involved in regulation of protein translation and RNA homeostasis, including *PELO*, *DDX6*, and *TUT7*, showed the strongest synthetic lethal effects with the SKI complex interacting protein-encoding gene *FOCAD*, which is frequently co-deleted with the tumor suppressor genes *CDKN2A/B* as collateral damage, among all examined gene-gene pairs. The identification of this interaction is consistent with recent studies identifying *PELO* and *TUT7* as synthetic lethal with *FOCAD* loss.^17,18,29^

To expand upon the results of our combinatorial enCas12a screens, we re-expressed *FOCAD* or an empty control vector in endogenously *FOCAD*-deleted MIAPACA2 cells to generate an isogenic cell line pair to investigate all possible genetic dependencies that are specific to *FOCAD* loss using a gRNA library targeting the whole genome. This orthogonal approach confirmed the enCas12a screen results and identified additional translation and RNA degradation factors *HBS1L*, *DIS3L2*, and *EDC4* as dependencies in *FOCAD*-deleted, but not *FOCAD* intact, cells. Among them, *HBS1L*, a gene encoding a ribosome rescue factor and obligate binding partner to the protein product of our enCas12a screen top hit *PELO*, demonstrated the strongest effect size. The interaction between *FOCAD* and *DIS3L2* was recently reproduced in literature but showed an overall weak effect compared to other interactions.^29^ The interaction between *FOCAD* and *EDC4* was not strongly reproduced by our internal in silico analysis tool TANDEM incorporating publicly available DepMap data, suggesting the effect may not be robust across aggregated data from many cell lines. The interactions between *FOCAD* and *HBS1L* and *PELO* were reproduced by our internal in silico analysis tool TANDEM incorporating publicly available DepMap data and are consistent with recent studies investigating synthetic lethal relationships with SKI complex inactivation^17,18^ .

Single-gene knockout of *HBS1L* in cell lines with *FOCAD* loss was lethal across multiple models, while cells with intact *FOCAD* did not depend on *HBS1L* for survival. The loss of viability observed upon *HBS1L* knockout in *FOCAD*-deleted cells could be rescued by expression of an edit-resistant *HBS1L* cDNA, demonstrating the on-target nature of the effect, or by re-expression of *FOCAD*, demonstrating the specificity of the context. FOCAD re-expression also resulted in corresponding stabilization of SKI complex protein SKIV2L, consistent with the proposed role of FOCAD in stabilizing the multiprotein SKI complex^13,17^ . Further, titration of HBS1L protein level using targeted protein degradation (HBS1L-dTAG fusion protein covering endogenous *HBS1L* knockout) showed a dose-dependent effect on viability of *FOCAD*-deleted MIAPACA2 cells, providing further pharmacological validation of the interaction. Inactivation of HBS1L led to translational arrest and induction of the unfolded protein response, consistent with the role of HBS1L in rescue of stalled ribosomes. *In vivo*, growth of xenograft tumors of *FOCAD*-deleted MIAPACA2 cells was completely abrogated by *HBS1L* depletion, demonstrating the robustness of this dependency.

The HBS1L/PELO complex is responsible for rescue of ribosomes that are stalled on aberrant mRNAs. These ribosomes can then continue to translate proteins needed for essential cellular function. To ensure the aberrant mRNA is not re-translated, it is cleaved by an endonuclease and the fragments are degraded by the exosome, in conjunction with the FOCAD-stabilized SKI complex, from the 3’→5’ direction and by XRN1 from the 5’→3’ direction (**Fig. 6A**). We propose a model in which tumor cells that have lost *FOCAD*, frequently through collateral damage from chromosomal deletion of the nearby *CDKN2A/B* tumor suppressor genes on chromosome 9p, lose the ability to degrade the 5’ end of aberrant mRNAs through destabilization of the SKI complex, exemplified by the reciprocal stabilization of SKI complex member SKIV2L when FOCAD expression is altered. These mRNAs then build up, and ribosomes can begin translation of them again and become stalled due to a lack of stop codon (**Fig. 6B**). These stalled ribosomes are dependent on HBS1L for rescue and recycling, which underlies the hyperdependence of *FOCAD*-deleted cells on HBS1L. Without HBS1L performing ribosome rescue, the increased bulk of aberrant mRNAs leads to ribosome sequestration through repeated and unrescued stalls, and the improperly synthesized partial proteins made by stalled ribosomes activates the unfolded protein response and leads to cell death (**Fig. 6B**).

**Figure 6.**
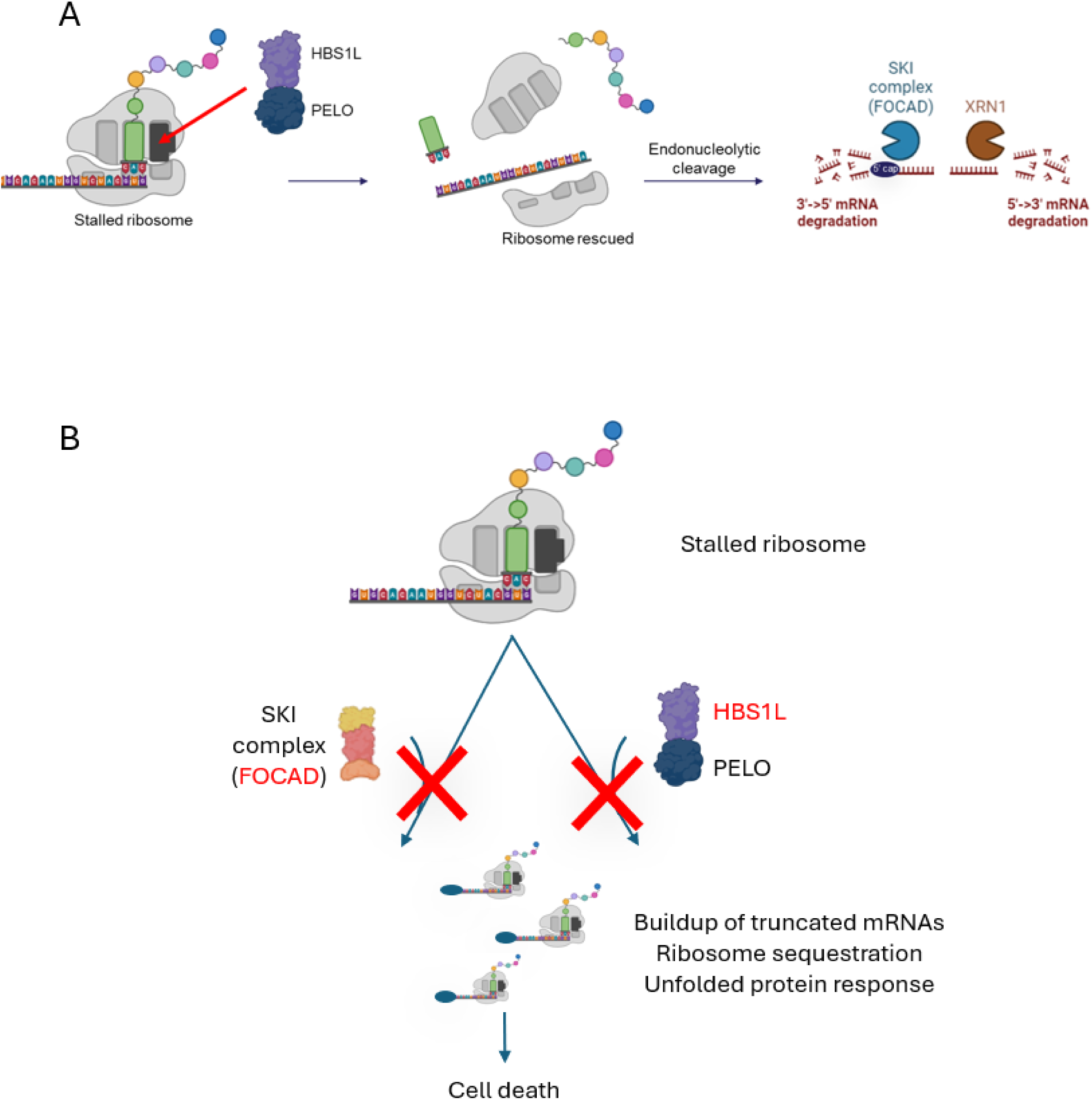
HBS1L/PELO and FOCAD are required for ribosome rescue and mRNA homeostasis. A) Model of the role of the HBS1L/PELO complex in ribosome rescue and subsequent role of FOCAD and the SKI complex in the degradation of aberrant mRNAs. B) Therapeutic hypothesis for the synthetic lethal interaction between HBS1L/PELO and FOCAD/SKI complex. Images created with BioRender (BioRender.com).

The *MTAP* gene sits between *FOCAD* and the *CDKN2A/B* tumor suppressor genes on chromosome 9p21. The majority of *FOCAD*-deleted tumors show co-deletion of both *CDKN2A/B* and *MTAP*, and conversely ∼1/3 of *MTAP*-deleted tumors also have *FOCAD* deletion (**Supplementary Fig. S9**). The synthetic lethal interaction between HBS1L and FOCAD provides an attractive novel opportunity for therapy that could be impactful for the ∼50,000 new cancer cases annually in the USA that have *FOCAD* loss, including ∼7.5% of NSCLC, and the strength of this relationship merits further development.

## Supporting information

EnCas12a library

Cas9 genome wide library

EnCas12a screen results

FOCAD context screen results

HBS1L and PELO dependencies

## ACKNOWLEDGEMENTS

We thank our partners at Pharmaron Inc (Beijing, China), ChemPartner (Shanghai, China), and BioMetas (Shanghai, China) for their support of our studies. The studies in this body of work were funded by Tango Therapeutics.

