## Supplementary Materials for "Tumor suppressor collateral damage screens reveal mRNA homeostasis protein HBS1L as a novel vulnerability in ch9p21 driven FOCAD deleted cancer"

**SUPPLEMENTAL MATERIALS for**

### Unbiased CRISPR screens identify synthetic lethal relationship between ribosome rescue factor HBS1L and mRNA homeostasis regular FOCAD

Hongxiang Zhang, Matthew R. Tonini, Lauren Catherine M. Martires, Charlotte B. Pratt, Helena N. Jenkins, Eden N. Gordon, Shanchuan Zhao, Ashley H. Choi, Samuel R. Meier, Tenzing Khendu, Shangtao Liu, Binzhang Shen, Hannah Stowe, Katerina Pashiardis, Xuewen Pan, Madhavi Bandi, Minjie Zhang, Yi Yu, Chengyin Min, Alan Huang, Jannik N. Anderson, Hilary E. Nicholson, Teng Teng

#### SUPPLEMENTARY METHODS

##### Combinatorial CRISPR gene interaction score analysis

FASTQ files were first mapped to the reference file of guide sequences. After mismatches and low-quality reads were removed, only reads perfectly aligned to all three sgRNA sequences at intended positions (**Fig. 1A**) were included in further analysis. As the sgRNAs at the second and third position were designed to target the same gene in druggable library, they were analyzed as a combined single sgRNA which paired with the sgRNA at the first position targeting the co-deleted context gene. A pseudo-count of 1 was added to all sgRNA pair counts in each sample to calculate the logarithmic fold change (log2FC) of each sgRNA guide pair count in endpoint relative to the plasmid sample. The sgRNA guide pair counts were normalized by the ratio between median count of non-targeting control (NTC) guide pairs (NTC-NTC) of the endpoint sample to the median count of NTC-NTC in the plasmid sample.  The normalized Log2FC for each sgRNA pair of each endpoint sample represents the observed effect of the sgRNA pair in the experiment condition.  The normalization assumes that NTC-NTC have no effect or association with the cancer cell survival, so that the median of Log2FC of NTC-NTC guide pairs equal to 0 in all samples. The normalized log2FC of guide pair of sgRNA $ij$ is the observed effect ($O_{ij}$) of the guide pair:

$$O_{ij} = \log_2\left(\frac{Ne_{ij}}{Np_{ij}}\right) - \log_2\left(\frac{median(Ne_{cc})}{median(Np_{cc})}\right) \quad (1)$$

Where $Ne_{ij}$ is the read count at the endpoint; $Ne_{ij}$ is the read count in plasmid of sgRNA pair $ij$; $Ne_{cc}$ and $Np_{cc}$ are the read count of all the NTC-NTC guide pairs at endpoint and in the plasmid respectively.

The single guide effect ($S_i$) of sgRNA $i$ was estimated by the mean observed effect of sgRNA pairs where the sgRNA was paired with a NTC guide ( $j \in C$). Similarly, $S_j$ was estimated.

$S_i$ = mean($O_{ij}$) where $j \in C$

$S_j$ = mean($O_{ij}$) where $i \in C$ (2)

For each sub-library of sgRNA pairs with the same context gene sgRNA $i$ at the first position, the observed effects were fitted by a quadratic model with the effect of the single sgRNA targeting druggable library gene ($Sj$) as predictor variable and the effect of the single sgRNA targeting the context gene ($Si$) as intercept. The Expected effect of each sgRNA pair ($E_{ij}$) was predicted based on the additive quadratic model of $Si$ and $Sj$.[22].

$$E_{ij} = \beta_1 S_j + \beta_2 S_j^2 + S_i \quad (3)$$

Guide pairs interaction effects (GIs) were residuals of the model, calculated by subtracting the expected sgRNA pair effects predicted by the quadratic model from the observed sgRNA pair effects.

$$GI_{ij} = O_{ij} - E_{ij} \quad (4)$$

As the variance of GIs getting greater when the expected effect size of the guide pair becomes more lethal, the standard deviation of GIs was corrected by dispersion adjustment[23]. The sgRNA pairs were ranked by the expected effect and the standard deviation of GIs was estimated in each bin of 1000 sgRNA pairs with similar rankings of expected effect.

$$\mathrm{Lsd}(GI_{ij}) = \sqrt{\frac{\sum(GI_{ik} - \overline{GI_{iw}})^2}{w-1}} \quad (5)$$

Where k=1,2,..$w$ in ascending ranking of guides in | $E_{ij}$ - $E_{ik}$ | defining the bin; $GI_{iw}$ is the sample mean of GI in the bin; Lsd($GI_{ij}$) is the local standard deviation of the bin surrounding $GI_{ij}$ ; $w$ is the number of nearby guides in the bin ($w$=1000).

$$aGI_{ij} = \frac{GI_{ij}}{Lsd(GI_{ij})} \quad (6)$$

The GI values were adjusted  by the local standard deviation.  Then adjusted GI values (aGI) were further converted to z-scores (zGI) by its mean and standard deviation.

$$zGI_{ij} = \frac{aGI_{ij} - \overline{aGI_i}}{sd(aGI_i)} \quad (7)$$

Finally, the average of zGI across all sgRNA pairs targeting co-deleted gene A and druggable library gene B were defined as the gene pair score ($GPS_{AB}$).

$$GPS_{AB} = \frac{\sum zGI_{ij}}{m} \ (i \in \text{guides targeting A}, \ j \in \text{guides targeting B}) \quad (8)$$

where $m$ is the number of guide pairs targeting A and B. The p-value of $GPS_{AB}$ was derived using the Stouffer's method where zGI of different sgRNA $ij$ guides pairs are viewed as independent test statistics[24]. Alternative meta-analysis methods, such as Lipták's weighted z-test or rank-based methods, could also be applied to combine the guide-pair statistics for further study. However, in our study, Stouffer's method provides a robust and powerful test for combining statistics across guide-pairs for gene-gene interaction analysis. Finally, the threshold levels of significance were adjusted for multiple tests by Bonferroni correction ($0.05/(5137*20) = 5x10^{-7}$).

**Table S1. Compounds**

| Compound | Vendor | Catalog No. |
|---|---|---|
| Crystal violet | Sigma Aldrich | V5265 |
| Doxycycline | R&D Systems | 4090 |
| dTAGv1 | R&D Systems | 6914 |

**Table S2. Cell lines, culture conditions and reagents, and engineering reagents**

| Cell Line/Reagent | Vendor | Catalog No. | Complete Medium |
|---|---|---|---|
| MIAPACA2 | ATCC | CRL-1420 | DMEM+10% FBS |
| A549 | ATCC | CCL-185 | DMEM+10% FBS |
| T24 | ATCC | HTB-4 | DMEM+10% FBS |
| MDAMB231 | ATCC | CRM-HTB-26 | DMEM+10% FBS |
| CFPAC1 | ATCC | CRL-1918 | IMDM+10%FBS |
| LN229 | ATCC | CRL-2611 | DMEM+5% FBS |
| ACHN | ATCC | CRL-1611 | EMEM+10%FBS |
| LN18 | ATCC | CRL-2610 | DMEM+5% FBS |
| Lenti-X | Takara Bio | 632180 | DMEM+10% FBS |
| HeLa | Abcam | ab255928 | EMEM+10%FBS |
| HeLa FOCAD knockout | Abcam | ab265627 | EMEM+10%FBS |
| DMEM | Gibco | 11965084 | |
| EMEM | Gibco | 11095072 | |
| IMDM | Gibco | 12440046 | |

| | | |
|---|---|---|
| FBS | GeminiBio | 110106 |
| MycoAlert Detection Kit | Lonza | LT07-318 |
| Lentiviral packaging mix | Cellecta | CPCP-K2A |
| Lipofectamine 3000 | Thermo Fisher Scientific | L3000015 |

**Table S3. Antibodies**

| **Target** | **Vendor** | **Catalog No.** | **Dilution** |
|---|---|---|---|
| PELO | Abcam | ab154335 | 1:1000 |
| HBS1L | Proteintech | 10359-1-AP | 1:1000 |
| IFRD1 | Abcam | Ab137633 | 1:1000 |
| CHOP | Cell Signaling | 5554 | 1:1000 |
| TRIB3 | Cell Signaling | 43043 | 1:1000 |
| GADD34 | Cell Signaling | 41222 | 1:1000 |
| HA tag | Cell Signaling | 3724 | 1:1000 |
| Puromycin | Sigma Aldrich | MABE343 | 1:1000 |
| β-actin | Cell Signaling | 3700 | 1:1000 |
| Vinculin | Cell Signaling | 13901 | 1:10,000 |

| | | | |
|---|---|---|---|
| Goat anti-mouse | ProteinSimple | 042-205 | N/A |
| Goat anti-rabbit | ProteinSimple | 042-206 | N/A |
| Goat anti-rabbit IR680 | LI-COR | 926-68071 | 1:15000 |
| Goat anti-mouse | LI-COR | 926-32210 | 1:15000 |

**Table S4. DNA sequences used in plasmids**

| Plasmid | Description | Sequence |
|---|---|---|
| pTG59, pTG760 | Intron-targeting control (sgITC) | GTACATGAAAAGGCTCTAGG |
| pTG2903 | sgHBS1L#1 | ATGCCTTGATCACATGAGAG |
| pTG2904 | sgHBS1L#2 | TTGTTTAAGGGGAGTAACCA |
| pTG3130, pTG3145, pTG2859 | HBS1L cDNA | GCCCGGCATCGGAATGTTCGAGGCTATAACTACGAT GAAGATTTTGAAGATGATGATCTCTACGGCCAGTCT GTAGAGGATGATTATTGTATTTCGCCGTCAACAGCT GCTCAGTTTATTTATTCACGGCGTGACAAACCTTCCG TTGAGCCTGTGGAAGAATATGATTATGAAGATCTGA AAGAATCTTCCAATTCTGTTTCAAACCATCAGCTCAG TGGATTTGATCAAGCTCGTCTTTATTCCTGTCTCGAC CATATGAGGGAAGTACTTGGAGATGCTGTGCCAGAT GAAATATTAATTGAAGCAGTTCTGAAGAACAAGTTT GATGTGCAGAAGGCTTTGTCAGGGGTTCTGGAACAA GATAGAGTGCAGAGTTTGAAGGACAAGAATGAGGC |

| | | |
|---|---|---|
| | | AACAGTATCTACAGGAAAGATAGCAAAAGGAAAAC CAGTAGATTCCCAGACATCGCGAAGTGAATCTGAAA TTGTGCCAAAAGTTGCTAAAATGACTGTATCTGGAA AGAAGCAAACTATGGGATTTGAAGTGCCTGGAGTAT CTTCTGAAGAAAATGGTCATAGTTTCCACACACCTC AAAAAGGACCGCCCATTGAAGATGCCATTGCTTCTT CCGATGTTCTTGAGACTGCTTCTAAATCTGCTAATCC ACCCCACACGATTCAAGCATCAGAAGAGCAGAGTTC AACCCCAGCACCGGTGAAAAAGTCTGGCAAGCTGA GGCAGCAAATAGATGTGAAGGCGGAACTGGAGAAG CGGCAAGGAGGGAAGCAGCTACTCAACTTAGTGGTC ATTGGTCATGTTGATGCTGGGAAAAGTACTCTGATG GGCCATATGCTTTATCTTCTGGGTAATATAAACAAA AGAACTATGCATAAGTATGAACAGGAGTCTAAAAA GGCTGGCAAAGCTTCGTTTGCATATGCATGGGTCTT GGATGAAACTGGCGAAGAAAGGGAGAGAGGCGTCA CAATGGATGTTGGTATGACAAAGTTTGAAACCACAA CCAAAGTTATTACATTAATGGATGCTCCAGGCCATA AGGACTTCATTCCAAATATGATTACAGGAGCAGCCC AGGCGGATGTAGCTGTTTTAGTTGTAGATGCCAGCA GGGGAGAGTTTGAAGCTGGATTTGAGACTGGAGGAC AAACACGAGAGCATGGACTCTTGGTCCGTTCTCTGG GAGTGACGCAGCTTGCAGTTGCAGTTAATAAAATGG |

| | | |
|---|---|---|
| | | ATCAGGTTAATTGGCAACAAGAAAGGTTTCAAGAGA TTACTGGAAAACTTGGGCACTTTCTTAAGCAAGCAG GTTTTAAGGAGAGTGATGTAGGTTTTATTCCTACAA GTGGTCTCAGTGGTGAAAATCTAATCACAAGATCTC AGTCAAGTGAACTCACAAAATGGTATAAAGGACTAT GTTTATTAGAACAAATTGATTCCTTTAAGCCTCCCCA GCGATCTATTGACAAACCTTTTAGATTATGTGTGTCC GATGTTTTCAAAGATCAAGGATCTGGATTTTGCATA ACTGGTAAAATAGAAGCTGGTTATATCCAAACTGGT GACCGACTACTGGCAATGCCTCCTAATGAAACTTGT ACCGTGAAAGGAATCACTCTGCATGATGAACCTGTC GACTGGGCGGCAGCAGGCGATCATGTTAGTCTTACT TTGGTTGGGATGGATATCATCAAAATCAATGTTGGC TGCATATTTTGTGGCCCCAAAGTACCCATTAAAGCTT GCACTCGTTTCAGAGCCCGAATCCTCATCTTTAATAT TGAAATTCCTATCACTAAAGGATTTCCTGTGCTGTTA CACTACCAAACTGTCAGTGAACCCGCCGTTATTAAA CGATTGATTAGTGTCTTAAACAAAAGCACGGGTGAA GTCACAAAGAAAAAGCCTAAGTTTTTGACTAAAGGC CAGAATGCATTGGTAGAGCTACAGACACAAAGACC AATAGCTCTTGAGCTATATAAAGACTTTAAAGAGCT GGGGAGGTTCATGCTACGTTACGGTGGTTCTACAAT AGCTGCTGGTGTTGTCACTGAGATAAAAGAATGA |

| | | |
|---|---|---|
| pTG3033 | FOCAD | ATGTCAGATGATATCAGGAAAAGGTTTGAATTTCCA<br>AATTCTCTTATCCAATCACAGGCTGTGGGTCATCTTA<br>TTGCTGCAGTCCTAAAGGAAAATGGTTTTTCAGAAA<br>AGATTCACCAATCTACAAATCAGACTCCTGCTTTGA<br>ACTTGCTGTGGGAGAAGTGTTGCAGTGACAATGTAG<br>TGGTTCGAACAGCCTGCTGTGAAGGTCTGGTGGCAC<br>TCGTTGCTCAGGATCATGCAGAGTTCAGCTATGTTCT<br>CAATGGGATACTCAACTTGATTCCATCAACCAGAAA<br>TACACATGGCTTGATAAAAGCCATTATGCACTTACT<br>ACAAATGCAAGCTCTTAAGGAAGGACAAGGTGGGG<br>AAAAGAATATTCAGAGTATATATACCATTAGAAATC<br>ATCCTCATCCTTTGATAACTGTGCTTGAACACAGACC<br>TGATTGCTGGCCAGTGTTTTTGCAGCAGCTGACAGC<br>GTTTTTCCAGCAGTGCCCTGAAAGGTTAGAAGTTTC<br>ATGCATTCAAATAATGGCACCATTTCTGTGGTATCTG<br>TATTGTGAACCATCTCAGTTACAAGAATATGCTAAA<br>CTCCGACTAGCCCTGCTGAAAGTCTTACTTCAACCCC<br>AGGTTCTTTGTGACAAAGATCAACCATCAATACTGG<br>AACAGCAGATACTTCAACTGTGTTGTGACATAGTTC<br>CATGTTTGCAGGTAAAAGATTTGATACAGACAACAG<br>AGGCGATGATGTTTATTGAGGAAGTATGTTTAAGCC<br>TTTTGCGTCATCCTGTTTTCTGGAAAATTCAGCTTAC<br>CCAGATGAGTCTTCAGCTGCTGTGTGTCAGTGAAGT |

| | | |
|---|---|---|
| | | CAGCTTAAAGATAACTGGTGAATGTTCATCTTCAATT CACCTTTTAGAGCACAGTGTTGAACTTCTGAAGGAG GATTTTCCTGTTGAACTGGTCATAATTGGAATAGCTT TACTACTTCTACAGACTCCAGCAAGTCAGCAGAAGC CAATCTTAAATCTAGCTTTGAAGCTCCTCTCTGTTAC TGAGGATCAGAAAATCCCAAAGTCCTCTCTGCTGCT AGTGATGCCAATTCTGCAGATACTATCTTCTACTGCC TTGGAAGACTGTATATCTGTGGATGAAGAAGGTCCC TCTAGGCAGCAGTTGGCTCTAAACCTTTTGGAAATG ATACAGCAGGAATGTTACAGAGATGACCACCAAAA GCTCTCCTACAAGCTTGTGTGCCCTGTAACCAGTATG TATGGTACAATATTTACAGCCTGGAGGATTCTTGAA GTAATGACAGACTCGTCTGCTGCAAGTGACTGGTTG GCTTCAGTAGAGTCATTGCTTCCTATTACTGCTGTGA TCCCTGCGCCTGCCTTTCTTCTGCTGGCTCACCTCCTT GTTGAAGACAAAGGACAAAATCTTCACCAAATACTC AAGGTCACTACAGAATTAGCCCAAGCAGATTCCTCC CAGGTGCCAAATCTGATTCCAGTTTTGATGTTCAAAT TGGGAAGACCACTGGAACCTATATTATATAATGATA TATTGTATACTTTACCTAAGCTTGGTGTTCACAAGGT GTGTATAGGACAAATTCTACGAATAATACAACTACT TGGAACCACACCACGACTAAGAGCTGTCACTTTGCG CTTGCTGACATCTTTGTGGGAAAAGCAGGACCGAGT |

| | | |
|---|---|---|
| | | CTATCCTGAACTGCAGCGTTTCATGGCTGTGTCTGAT GTACCTTCTCTTTCGGTGGGCAAGGAAGTCCAATGG GAGAAACTGATTGCAAAAGCAGCATCAATCAGAGA TATATGTAAGCAGAGGCCATATCAACATGGTGCAGA TATGTTGGCAGCTATTTCTCAAGTGTTGAATGAATGC ACCAAGCCTGATCAAGCTACTCCAGCAGCCTTGGTA TTACAGGGTCTTCATGCACTCTGTCAAGCTGAGGTTG TTTGCATTCGCTCCACTTGGAATGCTCTCTCTCCAAA GCTGAGTTGTGACACAAGACCTCTCATTCTGAAGAC ACTGAGTGAACTATTTTCTCTAGTTCCTTCCTTAACG GTCAATACAACTGAATATGAGAATTTTAAAGTTCAA GTCCTCAGCTTCCTCTGGACTCATACTCAAAACAAG GACCCAATTGTAGCAAATGCTGCATATAGATCCCTG GCCAACTTTAGTGCAGGAGAACACACCATTCTTCAT CTGCCTGAAAAGATAAGACCAGAAATTCCCATTCCT GAAGAGTTAGATGACGATGAAGATGTTGAGGATGTG GATCTTTCAGTTCCTGGCTCTTGCTATCTCAAACTGT TGTCACTCACTCCCCCTTTGGTTTTACCAGCTTTGGA GGAATTTTTTACATCACTTGTGAAGCAAGAAATGGT GAATATGCCTCGTGGGATATATCACTCTGCATTAAA AGGAGGTGCCCGCTCAGACCAAGGAAAGACTGTAG CAGGAATCCCCAATTTTATATTGAAAATGTATGAAA CAAACAAGCAACCAGGACTGAAACCTGGCCTTGCAG |

| | | |
|---|---|---|
| | | GTGGTATGTTATTTTGCTATGATGTTTCCATGTATCA<br>GAGTAAAGATGGAAAACCATTGAACAGACTGATGG<br>CCAGCAGAGGGCGAAGTTTCAAGCAGACTTCACTTG<br>CTCTTGTACATGAGGTTCATATCCAGCTTTCAGAGTG<br>GCACCGTGCAATTTTTCTTCCACAGGCCTGGCTTGCA<br>TACATGAATCGAGCTTATCATGCCATTTTACAGGGA<br>AGACTAGGAGAGCTGGAGTTGCAGTTAAAACATGG<br>AAAAGAAGAACCTGAGGAGGTGCAGTACAAAAAAA<br>GCACAGCCTGGCTCTGGGTTAGAGACATGCTGACTG<br>ATGAGATCACCAAGGCAGCTGCAAAGGAGAGTCCG<br>GTAGTGAAAGGCAATGCGCTGTTAGCTCTAAGCAGC<br>CTTGCTGTCGTCGTATCTAGACATGAAGCCAGCCTCT<br>CCTCAGACTCTGACGGGCTCCTGGAGGTTCAACCTA<br>ATTTCCTTTCAATGAAAGAGTGGGTTTCCATGGTACT<br>TGATACACTCTTGGTCATTGTGGATAGCCATTACCAA<br>CCCAGAGGGCAACTTCTCTCCTGGTTTTATTATAAGT<br>CCTATTCTGGTGAAAACACAGCTAGTGCCATTGCCC<br>GTTCTGCTGCCGCCACGGCTTTGTCTCTCCTTGTGCC<br>AGTTTTCATTATCTCTTGCAAAGAGAAGGTTGAGGA<br>AATCCTGAACATGCTGACTGCCAGGTTACCTGGGAA<br>ACCAAGTGCTGATGAGTCTCAAGCCGTGCAAATCCA<br>CATGGGCCTTGCTTTAGGGATGTTTCTCTCTCGCTTG<br>TGTGAAGAGAAACTCAGTGATATATCTGGCCAAGAG |

| | | |
|---|---|---|
| | | ATGAACCTTCTTCTGATGAAGTCGTTGGATGCCCTGG<br>AAAATTGCTGCTTTGACACTAGTCTTGAATACAACA<br>CGGGCTGTATATTGGGAGTTGGACTTGTTCTGTCCCT<br>CATGAGCCACAGCAGCCAAATGCAGTCCCGCGTTCA<br>CGTAGCAGCATTGCTCCGGAAGCTGTCTGCGCACGT<br>AGATGACAGCGGGAGCCAGAGCAGAACGTTTCAGG<br>AGGTCCTTGCCTACACACTTAGCTGTGTATGTACATC<br>AGCGTTCAGTGCTGGAATTATTGAGGCTACAGAGGC<br>TGAGGATGTTATGAACAAGCTTCGACTGTTAGTGGA<br>GAATAGCCAGCAGACTTCAGGTTTTGCCCTGGCTTT<br>AGGAAACATAGTTCATGGATTGTCTGTGTGTGGACA<br>TGGAAAAGCTGAAGACTTGGGCAGCAAACTACTCCC<br>TGCCTGGATCAGAATTGTTCTAACAGAGGGCACTCC<br>CACAATGCTTTGTCTGGCAGCTCTTCATGGCATGGTG<br>GCCTTGGTAGGCTCTGAAGGGGATGTAATGCAGCTG<br>AAATCAGAAGCCATCCAGACCTCTCATTTTCAAGGC<br>AGACTTAATGAAGTCATTAGAACCTTAACTCAGGTC<br>ATTAGTGTCTCTGGGGTGATTGGTCTCCAGTCAAATG<br>CAGTCTGGCTTCTTGGACATCTTCATCTATCTACTCT<br>ATCCTCAAGTCAAAGTAGAGCCTCTGTTCCTACTGA<br>CTATAGCTACTTGCCTGAAAGCAGTTTTATTGGAGC<br>AGCTATTGGCTTCTTCATTACAGGAGGAAAAAAAGG<br>TCCTGAATCTGTGCCTCCTTCCCTTCTTAAAGTAGTG |

| | | |
|---|---|---|
| | | ATGAAACCCATAGCAACTGTTGGAGAAAGCTACCAA<br>TATCCTCCTGTGAACTGGGCTGCACTTCTCTCTCCAC<br>TTATGAGGCTAAATTTTGGTGAAGAGATCCAGCAAC<br>TGTGCCTTGAAATTATGGTGACCCAGGCACAGTCAT<br>CCCAGAATGCAGCTGCACTATTGGGCTTGTGGGTGA<br>CACCACCACTGATCCACAGTCTGAGTCTGAATACCA<br>AGAGATATCTCCTGATATCTGCACCTCTGTGGATAA<br>AACACATCTCTGATGAACAGATCCTGGGTTTTGTTG<br>AAAATTTAATGGTGGCAGTTTTTAAAGCAGCTTCCC<br>CACTTGGAAGTCCTGAGCTATGCCCAAGTGCTTTAC<br>ACGGTCTGAGCCAGGCCATGAAACTGCCCAGCCCTG<br>CCCACCACCTCTGGAGTCTGCTCTCTGAAGCTACTGG<br>GAAAATTTTTGACCTCCTGCCAAATAAGATTCGGAG<br>AAAGGATCTAGAGCTGTATATCAGCATAGCAAAATG<br>CCTCTTAGAAATGACAGATGATGATGCCAATCGGAT<br>CGCCCAGGTTACTAAGAGCAACATAGAAAAAGCTGC<br>CTTTGTCAAACTGTACTTAGTCTCTCAAGGACGATTC<br>CCCTTGGTGAACCTGACCGATATGCTGAGCGTTGCT<br>GTGCAGCACCGTGAGAAAGAGGTGTTGGCCTGGATG<br>ATTCTGCACAGCTTATACCAGGCACGGATTGTGAGC<br>CATGCCAATACGGGCGTTTTGAAGAGAATGGAGTGG<br>CTCTTGGAACTGATGGGTTATATTAGAAATGTTGCTT<br>ACCAGTCAACATCCTTTCACAATACGGCTCTTGACA |

| | | |
|---|---|---|
| | | AGGCTTTGGACTTCTTCTTGCTGATATTTGCAACCGC AGTGGTTGCATGGGCTGACCACACTGCCCCTCTCCTC CTCGGCCTCAGTGCCAGTTGGTTGCCATGGCATCAG GAGAATGGCCCGGCTGGGCCAGTACCAAGCTTCCTT GGCAGGAGTCCAATGCACAGGGTCACTCTGCAGGAG GTTCTCACTCTCCTTCCCAATAGCATGGCTCTGCTGC TGCAGAAAGAGCCATGGAAGGAACAGACCCAGAAG TTCATTGACTGGCTATTCAGCATCATGGAAAGCCCT AAAGAAGCCCTCTCAGCACAGTCCAGGGATCTTTTG AAAGCCACCCTGCTGTCCTTGAGAGTTCTCCCAGAG TTTAAGAAGAAAGCTGTATGGACCAGAGCATATGGT TGGTGA |
| pTG3130 | FKBP12(F36V)-2XHA | ATGGGAGTGCAGGTGGAAACCATCTCCCCAGGAGAC GGGCGCACCTTCCCCAAGCGCGGCCAGACCTGCGTG GTGCACTACACCGGGATGCTTGAAGATGGAAAGAA AGTTGATTCCTCCCGGGACAGAAACAAGCCCTTTAA GTTTATGCTAGGCAAGCAGGAGGTGATCCGAGGCTG GGAAGAAGGGGTTGCCCAGATGAGTGTGGGTCAGA GAGCCAAACTGACTATATCTCCAGATTATGCCTATG GTGCCACTGGGCACCCAGGCATCATCCCACCACATG CCACTCTCGTCTTCGATGTGGAGCTTCTAAAACTGGA AGGCGGCTACCCCTACGACGTGCCCGACTACGCCGG CTATCCGTATGATGTCCCGGACTATGCA |

| | | |
|---|---|---|
| pTG3145 | FKBP12WT-2XHA | ATGGGAGTGCAGGTGGAAACCATCTCCCCAGGAGACGGGCGCACCTTCCCCAAGCGCGGCCAGACCTGCGTGGTGCACTACACCGGGATGCTTGAAGATGGAAAGAAATTTGATTCCTCCCGGGACAGAAACAAGCCCTTTAAGTTTATGCTAGGCAAGCAGGAGGTGATCCGAGGCTGGGAAGAAGGGGTTGCCCAGATGAGTGTGGGTCAGAGAGCCAAACTGACTATATCTCCAGATTATGCCTATGGTGCCACTGGGCACCCAGGCATCATCCCACCACATGCCACTCTCGTCTTCGATGTGGAGCTTCTAAAACTGGAAGGCGGCTACCCCTACGACGTGCCCGACTACGCCGGCTATCCGTATGATGTCCCGGACTATGCA |

**Table S5. qPCR probes**

| Gene | Supplier | Catalog No. |
|---|---|---|
| IFRD1 | Life Technologies | Hs00155477_m1 |
| CHOP/GADD153 | Life Technologies | Hs00358796_g1 |
| TRIB3 | Life Technologies | Hs01082394_m1 |

| | | |
|---|---|---|
| GADD34 | Life Technologies | Hs000169585_m1 |
| Human RPLPO-VIC/MGB | Life Technologies | 4326314E |

### SUPPLEMENTARY FIGURES

| ID | gRNA sequence | Reads_aligned | Indel % | Frameshift % |
|---|---|---|---|---|
| ACTA2_gRNA1 | CCATCCATTGTGGGACGTCCCAG | 4501 | 101% | 67% |
| ACTA2_gRNA2 | AGGGGTGCCTCCGTGAGCAGGGT | 50571 | 100% | 68% |
| **ACTA2_gRNA3** | TATCGGGTACTTCAGGGTCAGGA | 38175 | 99% | 72% |
| ACTA2_gRNA4 | TCACGCCCAATTGTCTCCCAGGC | 20387 | 101% | 64% |
| ATAD1_gRNA1 | TACGAAACCGTTCAAGTTCTGAC | 860 | 94% | 60% |
| ATAD1_gRNA2 | GATGATGTCATTACGGATCTGAA | 640 | 65% | 36% |
| **ATAD1_gRNA3** | TCGGTCAGTGTCGAAGGCTGAAG | 11555 | 100% | 73% |
| ATAD1_gRNA4 | ATCTATAAAGATGATGGATGGTT | 11180 | 101% | 71% |
| **DMRTA1_gRNA1** | GAAAGCGGGGGTCGCGGAACCAC | 11394 | 98% | 67% |
| DMRTA1_gRNA2 | ATTGGGAAACAAAGTATCGGGTC | 1062 | 81% | 52% |
| DMRTA1_gRNA3 | CAGATCAGAGGATGATAAGGACC | 1171 | 74% | 49% |
| DMRTA1_gRNA4 | GTGTCAGCGCTCAAGGGCCACAA | 49400 | 99% | 64% |
| ELAC1_gRNA1 | GTCGTGGAAAAGAAACGCCCAGG | 162190 | 24% | 10% |
| **ELAC1_gRNA2** | GCCAGATAAAGTCCCGAAGCCCT | 36651 | 90% | 64% |
| ELAC1_gRNA3 | CTACAGCAGATCAATGTCCTGCA | 300789 | 48% | 25% |
| ELAC1_gRNA4 | TGAGTCTAACAGGATAGTTCTTC | 348487 | 38% | 25% |
| **ELAVL2_gRNA1** | GCTTGGGTTATTAGCAAACTTTA | 94450 | 98% | 67% |
| ELAVL2_gRNA2 | CTCAAATCTACCATAGGCATATC | 17992 | 3% | 2% |
| ELAVL2_gRNA3 | GGAAGTCCGCTGACATATAAATT | 368 | 101% | 57% |
| ELAVL2_gRNA4 | AAGTCTCAATCCATTCAGGGTGT | 13 | 100% | 92% |
| FNDC3A_gRNA1 | GGACCCCCTCTGGTTGATGGTGG | 4501 | 99% | 64% |
| FNDC3A_gRNA2 | ATATCCATTCATACCGTTAGAAC | 528 | 121% | 69% |
| FNDC3A_gRNA3 | TCACCATAGAGTCCAGGCAGAAT | 443 | 97% | 65% |
| **FNDC3A_gRNA4** | TGTGAGCCCATGTAACACTGACA | 7578 | 97% | 67% |
| FOCAD_gRNA1 | CAGGGCATCCAACGACTTCATCA | 283 | 106% | 46% |
| **FOCAD_gRNA2** | AATCACCCCAACTAACCTCCAGG | 14939 | 101% | 72% |
| FOCAD_gRNA3 | CCATAGGTTCATATCCAGCTTTC | 10 | 100% | 30% |
| FOCAD_gRNA4 | GGTGCCAAATCTGATTCCAGTTT | 15 | 100% | 93% |
| **HACD4_gRNA1** | ACCCACGGGATACAGATTCAATG | 38484 | 100% | 55% |
| HACD4_gRNA2 | ACTCAATGCCAACATATATGTGC | 126032 | 62% | 40% |
| HACD4_gRNA3 | ACTGAGCCATGTCAAGACAGCAT | 1394 | 101% | 68% |
| HACD4_gRNA4 | TGCTATTGGACTTGTGATGCGAC | 125803 | 50% | 34% |
| **ITM2B_gRNA1** | GATCTTAACCTGGATAAGTGCTA | 14354 | 102% | 68% |
| ITM2B_gRNA2 | GGACCCAGATGATGTGGTACCAG | 3849 | 88% | 54% |
| ITM2B_gRNA3 | TAGGAGGAGCATACTTGTACAAA | 14113 | 95% | 54% |
| ITM2B_gRNA4 | AATGAGCCCTCTGCAGATGCCCC | 9778 | 95% | 66% |
| KLHL9_gRNA1 | GGTGGTATTGCAAGGCTTTGATC | 1594 | 48% | 30% |
| KLHL9_gRNA2 | AGACCAACCTTGTTCACCCCATG | 381 | 98% | 65% |
| **KLHL9_gRNA3** | GATTTCCACTGATGACACCACAG | 13350 | 100% | 60% |
| KLHL9_gRNA4 | CAGATTTGATCCTCGGTATAATA | 55 | 98% | 69% |

| ID | gRNA sequence | Reads_aligned | Indel % | Frameshift % |
|---|---|---|---|---|
| **MBD2_gRNA1** | GATTGAGAGGATCGTTTCGCAGT | 57474 | 102% | 66% |
| MBD2_gRNA2 | CAAACAACCGGTAACCAAAGTCA | 4075 | 99% | 57% |
| MBD2_gRNA3 | CCGACGGGAAAGGGACCGGCTCC | 106527 | 79% | 53% |
| MBD2_gRNA4 | TAGTCCAAGTGGTAAGAAGTTCA | 66 | 95% | 41% |
| ME2_gRNA1 | GATATGGCCGGAACACACTCATT | 4038 | 11% | 8% |
| ME2_gRNA2 | TAGGCACTCTTAAAAGACCCATT | 9 | 100% | 56% |
| ME2_gRNA3 | GTGCTGCAAGAAGACCTGCTAGA | 14499 | 97% | 55% |
| **ME2_gRNA4** | AATACCTTACCTTAACTAATAAA | 18976 | 99% | 70% |
| **MRO_gRNA1** | GCACTGGGGTCCCGAGCTCTTTC | 611 | 54% | 44% |
| MRO_gRNA2 | GTTGATGGGGACCGCAGGAGAAC | 6206 | 28% | 24% |
| MRO_gRNA3 | ACAGGGTCATACAGTCCATACAC | 4935 | 100% | 43% |
| MRO_gRNA4 | CTCCCGGCAAAGGCAGCCAATTG | 6281 | 72% | 36% |
| **PAPSS2_gRNA1** | AGGATCGTGAGAATGCCCGCAAA | 4067 | 100% | 66% |
| PAPSS2_gRNA2 | AGCCTCAGCTCGGACGTGGTCAA | 225 | 2% | 1% |
| PAPSS2_gRNA3 | AGCACACGCTCAGGAGTTTCAGG | 40 | 103% | 90% |
| PAPSS2_gRNA4 | ATGCGGGAGAAGGAGTACTTACA | 204 | 145% | 99% |
| RCBTB1_gRNA1 | GACTGAACTATAGTAACTGTCTA | 7 | 86% | 43% |
| **RCBTB1_gRNA2** | GTGCTAGAGTATGTGCGTAACCG | 7120 | 101% | 74% |
| RCBTB1_gRNA3 | AAAGCCTCAGTTACGGGAGTGGA | 99 | 66% | 45% |
| RCBTB1_gRNA4 | CATATTAAGAGGGTAGTTGGCAT | 2417 | 92% | 66% |
| RCBTB2_gRNA1 | CGTTGTAACCCCAGACATAGACC | 890 | 99% | 53% |
| RCBTB2_gRNA2 | GGGTTAGGTGACGTCCAGAGCAC | 2 | 150% | 50% |
| **RCBTB2_gRNA3** | GGATTATCGAGATTGCAGCCTGT | 2584 | 101% | 74% |
| RCBTB2_gRNA4 | AGGTCGCCTGTGGCTACGCACAC | 5655 | 99% | 69% |
| RNASEH2B_gRNA1 | GAAACACGTTATCCACCACAACT | 14 | 100% | 57% |
| RNASEH2B_gRNA2 | ATGCTGCCACAGTTTGATTAACC | 522 | 20% | 11% |
| **RNASEH2B_gRNA3** | CTTGTTCAATATGTGTCTACAGC | 350 | 96% | 58% |
| RNASEH2B_gRNA4 | TCTGGTGACCAAGCTTCCACTGA | 510 | 88% | 54% |
| **RNLS_gRNA1** | GACAACCCACCTGAGTCCTCAGC | 214162 | 100% | 80% |
| RNLS_gRNA2 | AGGCCTCTAAGCTCGCCTATTGA | 350 | 95% | 68% |
| RNLS_gRNA3 | ACTCCTTGCAACTAACTCACCGT | 1348 | 80% | 57% |
| RNLS_gRNA4 | GGTTGATCTGTGTCACACGATGT | 22916 | 99% | 78% |
| SETDB2_gRNA1 | CCTGCCTCACAGGAGCATCACAG | 617 | 100% | 59% |
| SETDB2_gRNA2 | CGAGCCAACTGAACATAGGTATT | 53146 | 43% | 27% |
| SETDB2_gRNA3 | ATAACTCACGTGGAGTGCTGAAG | 88356 | 63% | 38% |
| **SETDB2_gRNA4** | AACTGACAGCAAGGAATGCCAAA | 88356 | 97% | 62% |
| STAMBPL1_gRNA1 | GTGGTTGAAGGACTGCGATGTGT | 117 | 100% | 66% |
| **STAMBPL1_gRNA2** | CACACACCAGAACAATTCCTTGC | 12786 | 91% | 56% |
| STAMBPL1_gRNA3 | ACCTCGGGCTAACTCTTGCTTCT | 3080 | 97% | 60% |
| STAMBPL1_gRNA4 | ACTGATGGTGATATTACAACCAA | 6 | 100% | 67% |

**Supplementary Figure S1. EnCas12a gRNA editing efficiency evaluation for 20 context genes**

Editing efficiency evaluation of Encas12a gRNA for the 20 frequently co-deleted context genes. 4 gRNAs were evaluated per gene, the number of NGS sequencing reads aligned, the insertion and deletion reads percentage (INDEL%) and frameshift mutation rate were listed. The top gRNA for each gene is highlighted and selected for combinatorial library construction.

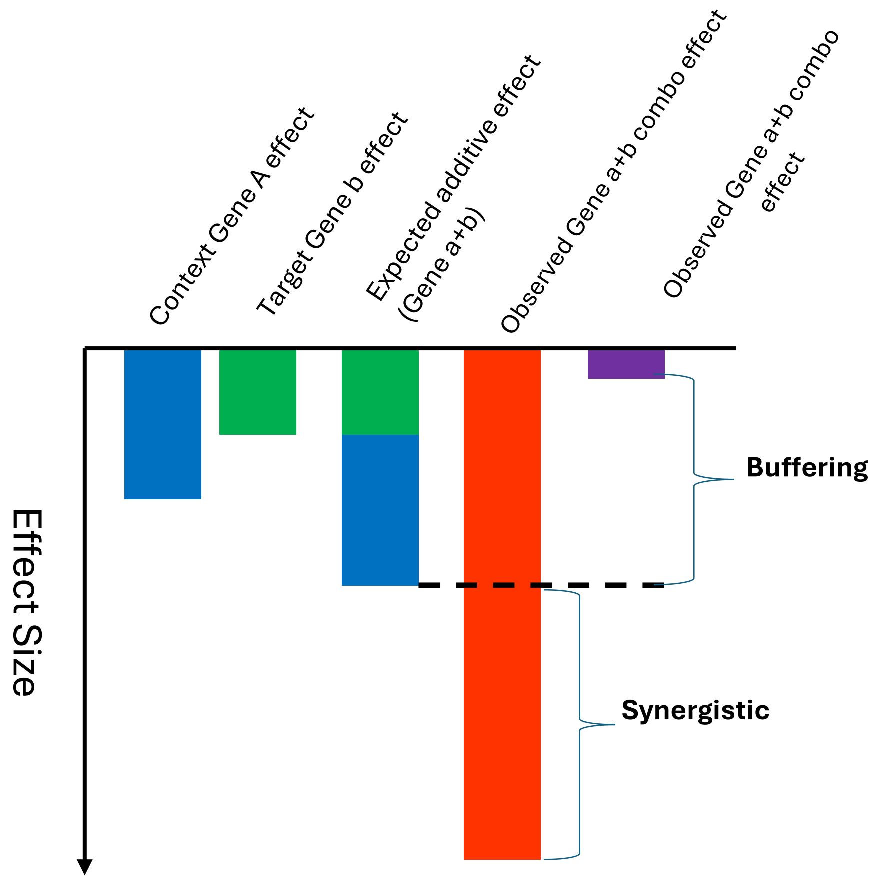

**Supplementary Figure S2. EnCas12a combinatorial screen synergistic effect calculation scheme**

Scheme for EnCas12a combinatorial library synergistic effect calculation (see methods for details). Single gene effect of both the context Gene A and target gene B can be estimated from the library result when paired with negative control gRNAs. Gene-gene synergistic interaction score can therefore be calculated based on the difference between the expected additive Gene A and B single knockout effect versus the observed Gene A and Gene B combinatorial knockout effect.

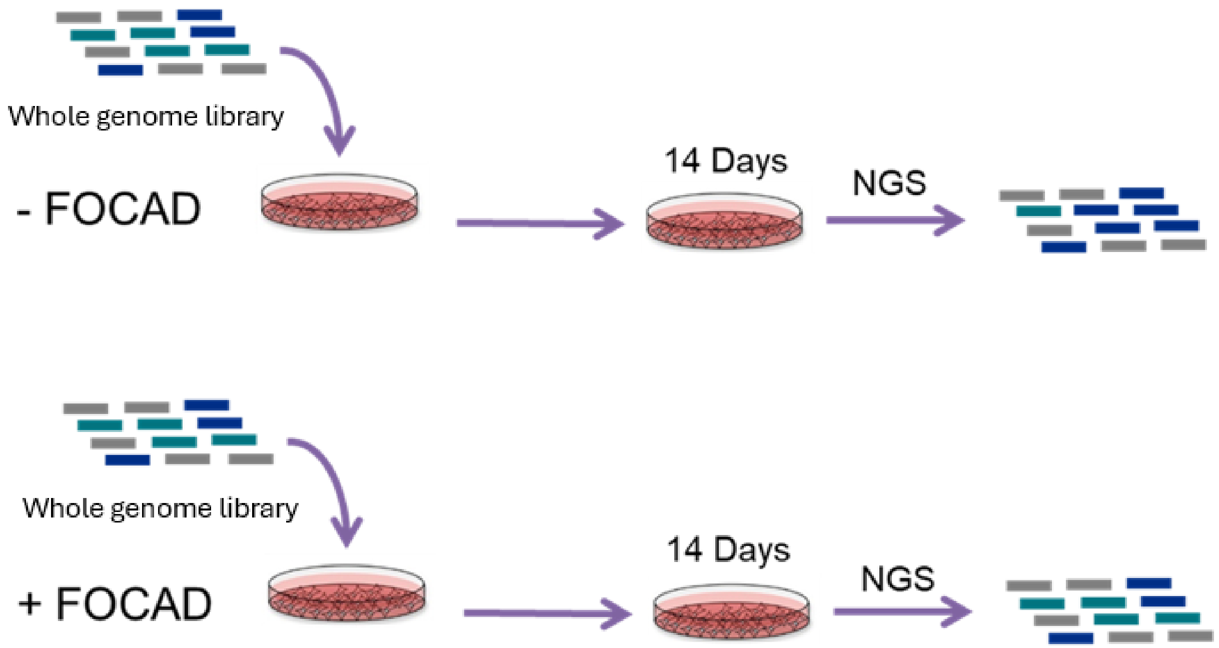

**Supplementary Figure S3. Whole genome SpCas9 screen scheme in engineered FOCAD isogenic pair cell lines (MIAPACA2)**

FOCAD-deleted MIAPAC2 Cas9 clone were infected with FOCAD expressing cDNA (+FOCAD) or empty vector (-FOCAD) and infected with whole genome library. 14 Days post puromycin selection, cell pellets were collected and submitted for NGS sequencing.

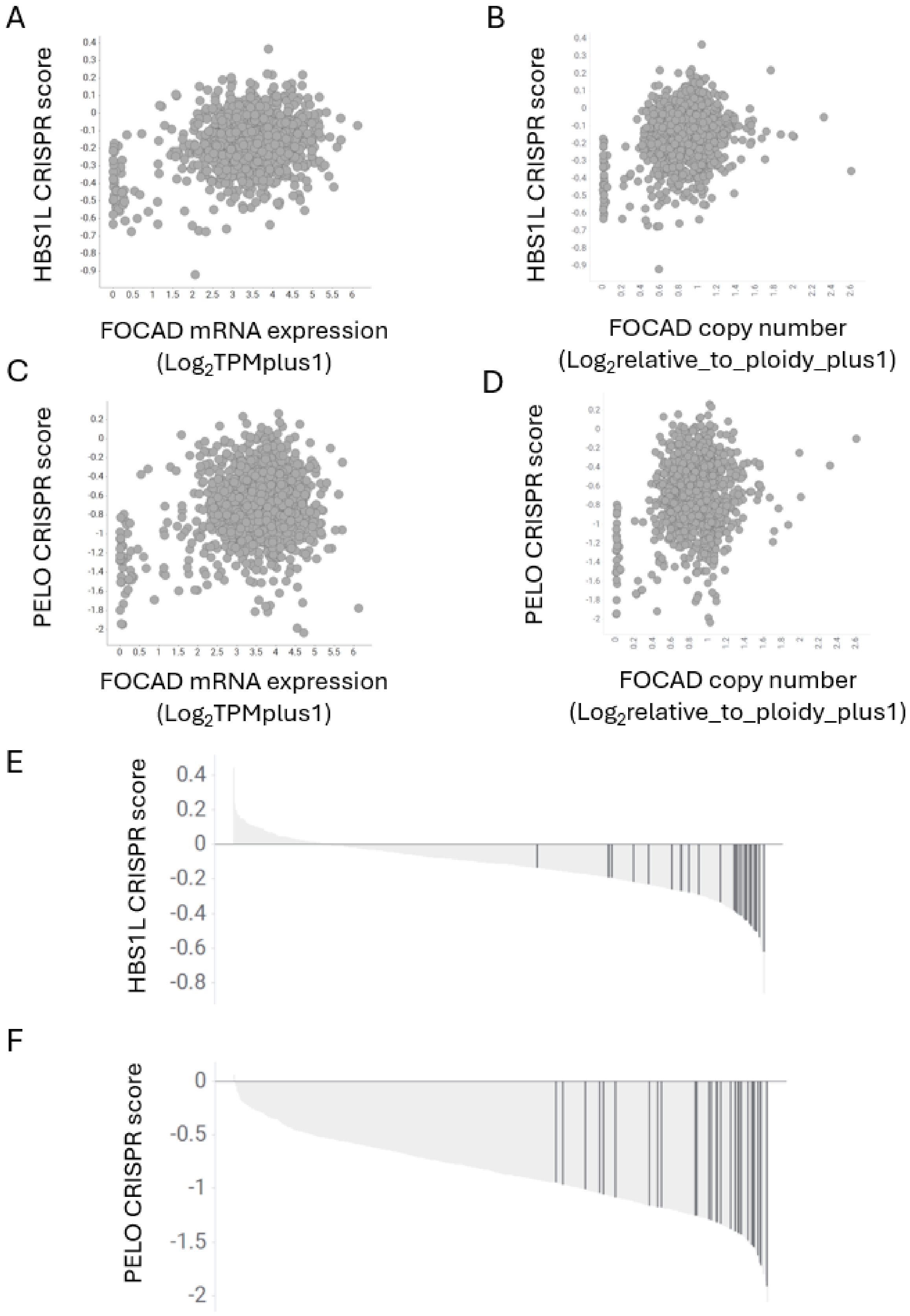
A
HBS1L CRISPR score
FOCAD mRNA expression
(Log2TPMplus1)
B
HBS1L CRISPR score
FOCAD copy number
(Log2relative_to_ploidy_plus1)
C
PELO CRISPR score
FOCAD mRNA expression
(Log2TPMplus1)
D
PELO CRISPR score
FOCAD copy number
(Log2relative_to_ploidy_plus1)
E
HBS1L CRISPR score
0.4
0.2
0
-0.2
-0.4
-0.6
-0.8
F
PELO CRISPR score
0
-0.5
-1
-1.5
-2

G

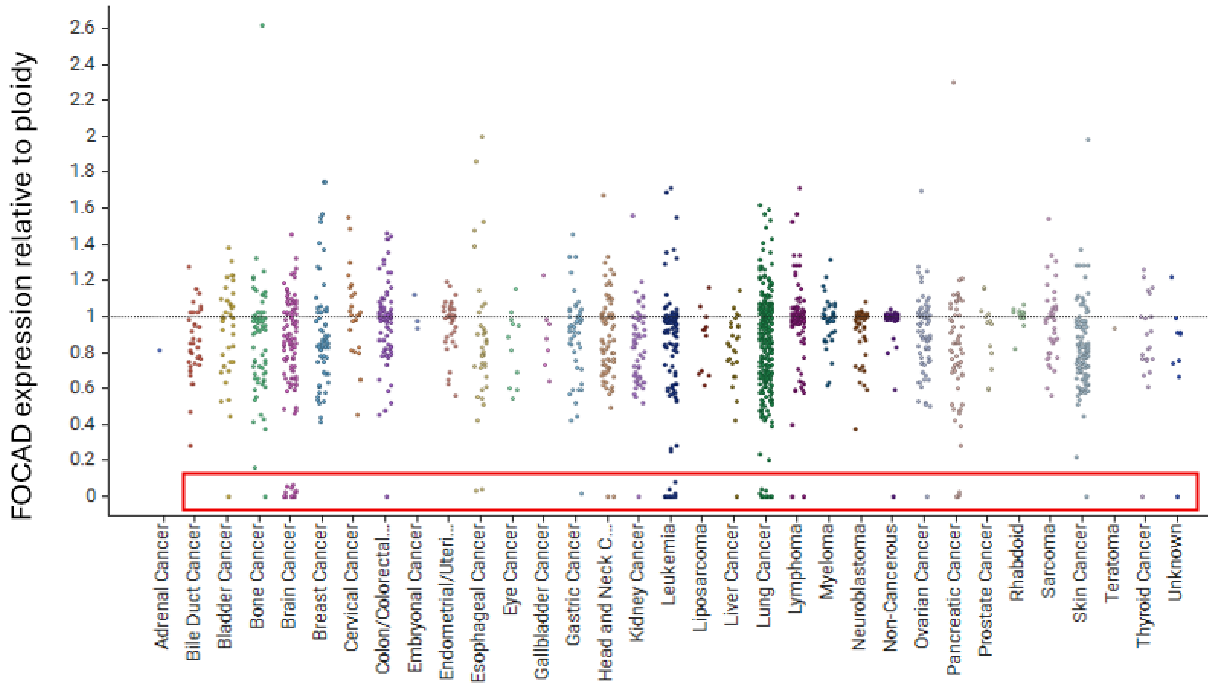
FOCAD expression relative to ploidy
2.6
2.4
2.2
2
1.8
1.6
1.4
1.2
1
0.8
0.6
0.4
0.2
0
Adrenal Cancer
Bile Duct Cancer
Bladder Cancer
Bone Cancer
Brain Cancer
Breast Cancer
Cervical Cancer
Colon/Colorectal...
Embryonal Cancer
Endometrial/Uteri...
Esophageal Cancer
Eye Cancer
Gallbladder Cancer
Gastric Cancer
Head and Neck C...
Kidney Cancer
Leukemia
Liposarcoma
Liver Cancer
Lung Cancer
Lymphoma
Myeloma
Neuroblastoma
Non-Cancerous
Ovarian Cancer
Pancreatic Cancer
Prostate Cancer
Rhabdoid
Sarcoma
Skin Cancer
Teratoma
Thyroid Cancer
Unknown

H

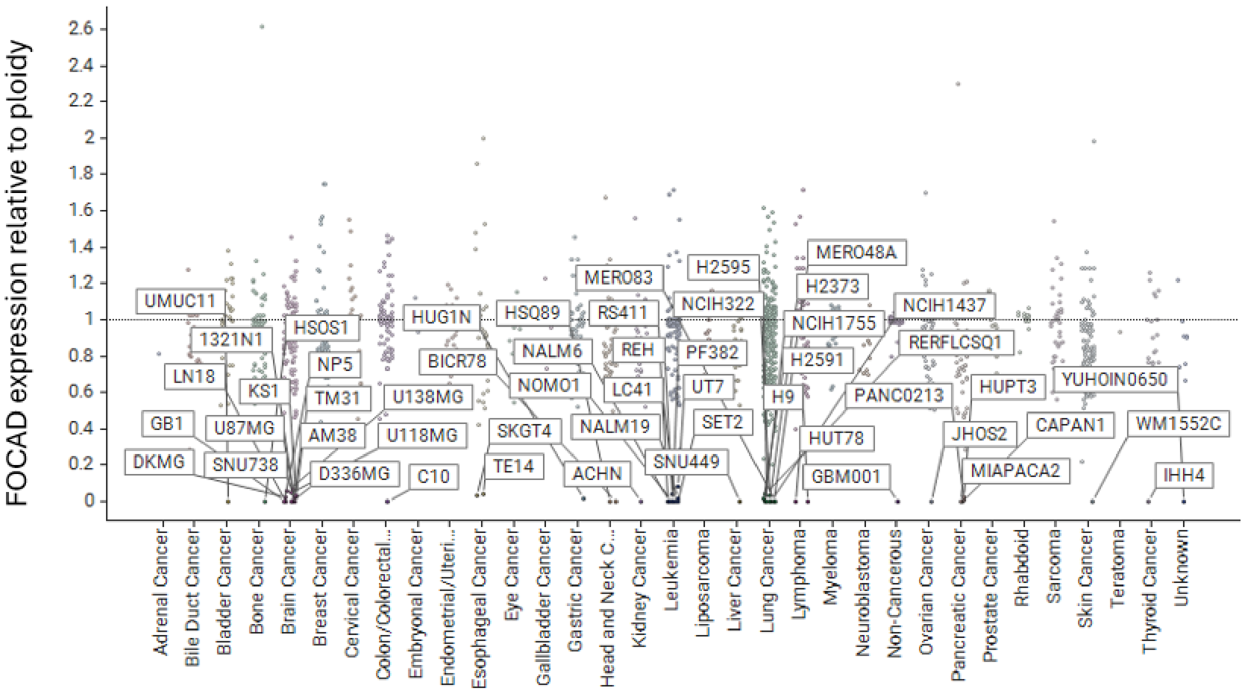
FOCAD expression relative to ploidy
2.6
2.4
2.2
2
1.8
1.6
1.4
1.2
1
0.8
0.6
0.4
0.2
0
UMUC11
1321N1
LN18
GB1
DKMG
U87MG
SNU738
KS1
HSOS1
NP5
TM31
AM38
D336MG
U138MG
U118MG
C10
HUG1N
BICR78
HSQ89
NALM6
NOMO1
SKGT4
TE14
ACHN
MER083
RS411
REH
LC41
NALM19
SNU449
H2595
NCIH322
PF382
UT7
SET2
H9
MERO48A
H2373
NCIH1755
H2591
HUT78
GBM001
NCIH1437
RERFLCSQ1
PANC0213
HUPT3
JHOS2
MIAPACA2
CAPAN1
YUHOIN0650
WM1552C
IHH4
Adrenal Cancer
Bile Duct Cancer
Bladder Cancer
Bone Cancer
Brain Cancer
Breast Cancer
Cervical Cancer
Colon/Colorectal...
Embryonal Cancer
Endometrial/Uteri...
Esophageal Cancer
Eye Cancer
Gallbladder Cancer
Gastric Cancer
Head and Neck C...
Kidney Cancer
Leukemia
Liposarcoma
Liver Cancer
Lung Cancer
Lymphoma
Myeloma
Neuroblastoma
Non-Cancerous
Ovarian Cancer
Pancreatic Cancer
Prostate Cancer
Rhabdoid
Sarcoma
Skin Cancer
Teratoma
Thyroid Cancer
Unknown

I

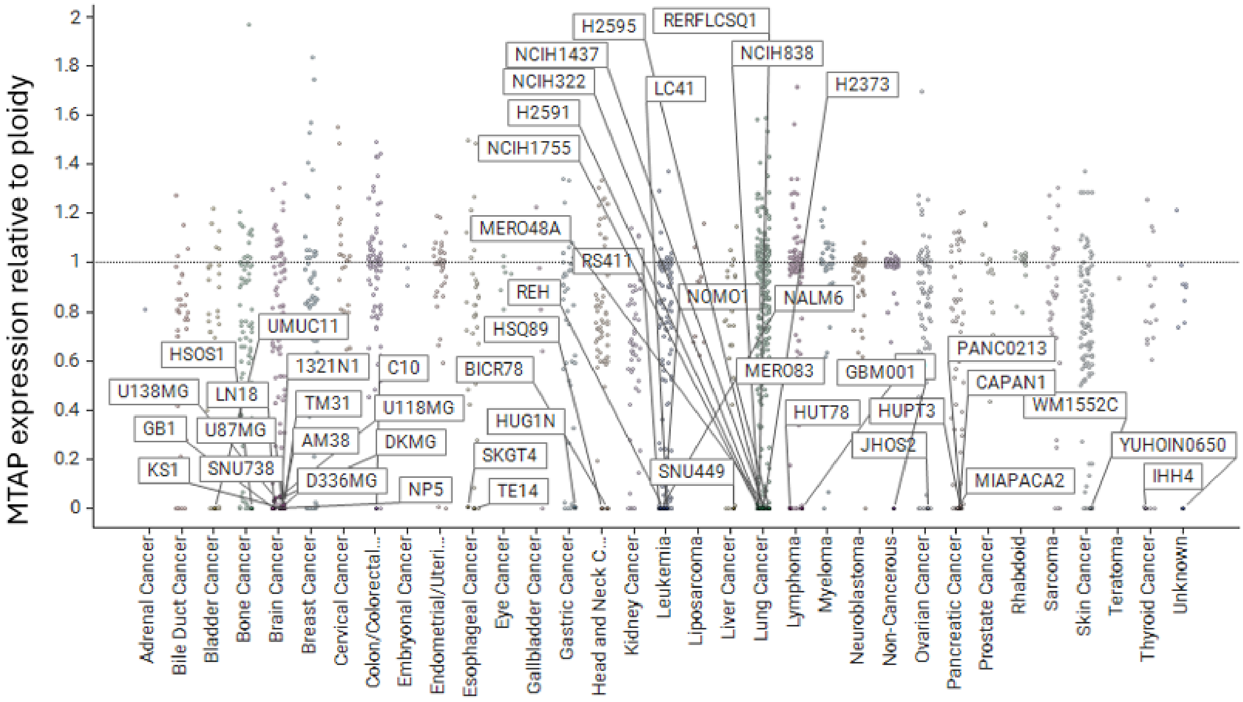

**Supplementary Figure S4. HBS1L shows the best correlation to FOCAD expression and copy number status in Achilles screen**

Correlation of HBS1L CRISPR score vs. FOCAD expression levels (A) and FOCAD copy number (B) in Achilles screen panel. Correlation of PELO CRISPR score vs. FOCAD expression levels (C) and FOCAD copy number (D) in Achilles screen panel. E) Waterfall plot of HBS1L CRISPR score across Achilles screen panel. FOCAD negative cell lines were highlighted. CRISPR scores by cell line can be found in Supplementary Dataset 5. F) Waterfall plot of PELO CRISPR score across Achilles screen panel. FOCAD negative cell lines were highlighted. (corresponding to Fig. 1E). CRISPR scores by cell line can be found in Supplementary Dataset 5. G, H) Annotation of cell lines having copy number deletion of FOCAD, boxed in red (G) and with cell line names (H). I) MTAP copy number status of FOCAD-deleted cell lines defined in panels G-H.

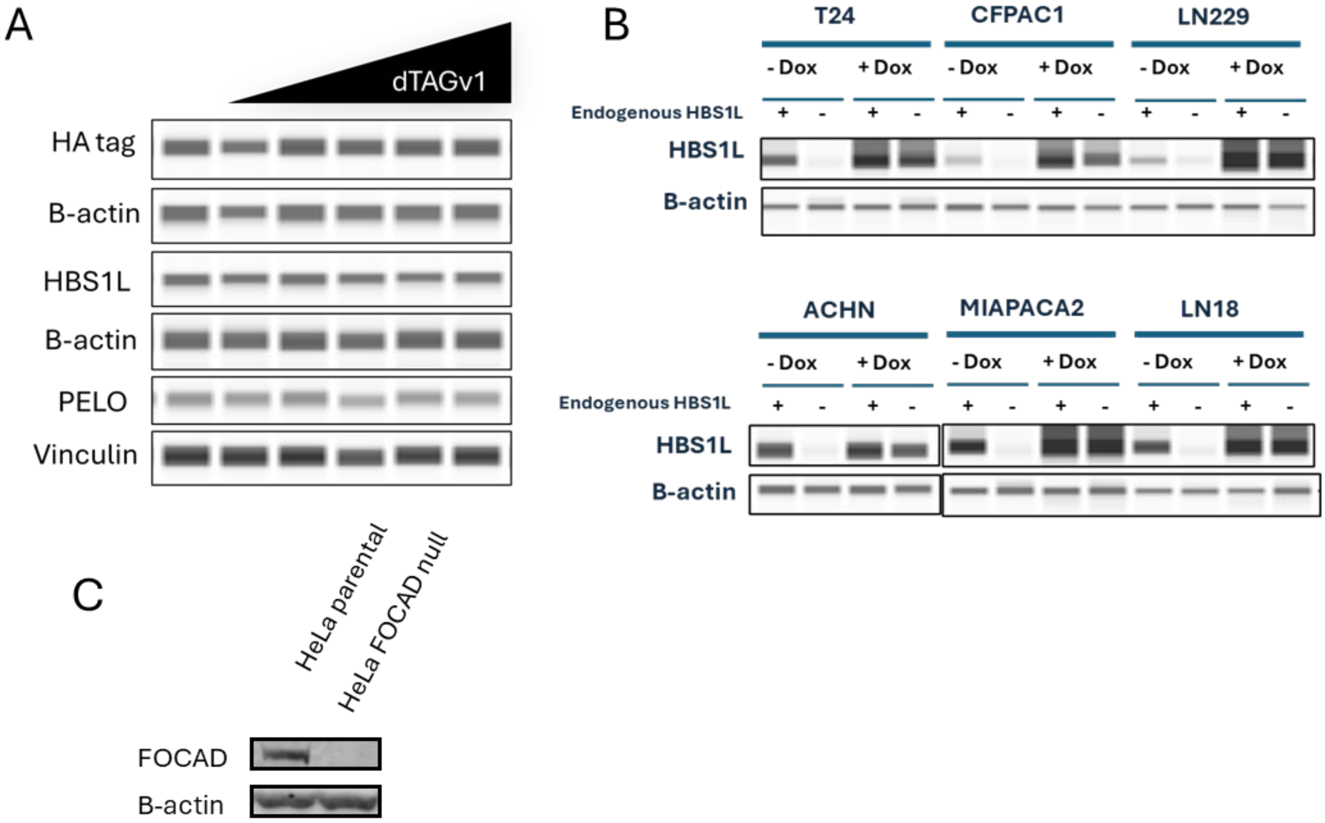

**Supplementary Figure S5. Characterization of protein expression in MIAPACA2-HBS1L-FKBP12WT cells, FOCAD-positive and FOCAD-negative minipanel of cell lines, and HeLa *FOCAD* isogenic cell lines**

A) Western blot of a clonal population of MIAPACA2 cells stably expressing *HBS1L-FKBP12WT* fusion cDNA and having endogenous knockout of *HBS1L* treated with 0, 0.5, 1, 5, 20, or 50 nM dTAGv1 for 14 days (corresponding to endpoint for colony formation assay in Fig. 2F). B) Western blot of HBS1L in indicated cell lines expressing DOX-inducible *HBS1L* cDNA covering endogenous *HBS1L* knockout or control sgRNA (indicated by “endogenous HBS1L”) (corresponding to cell lines used in colony formation assays in Fig. 3A). C) Western blot of FOCAD in HeLa parental and *FOCAD* knockout isogenic lines (corresponding to cell line used for colony formation assay in Fig. 3E)

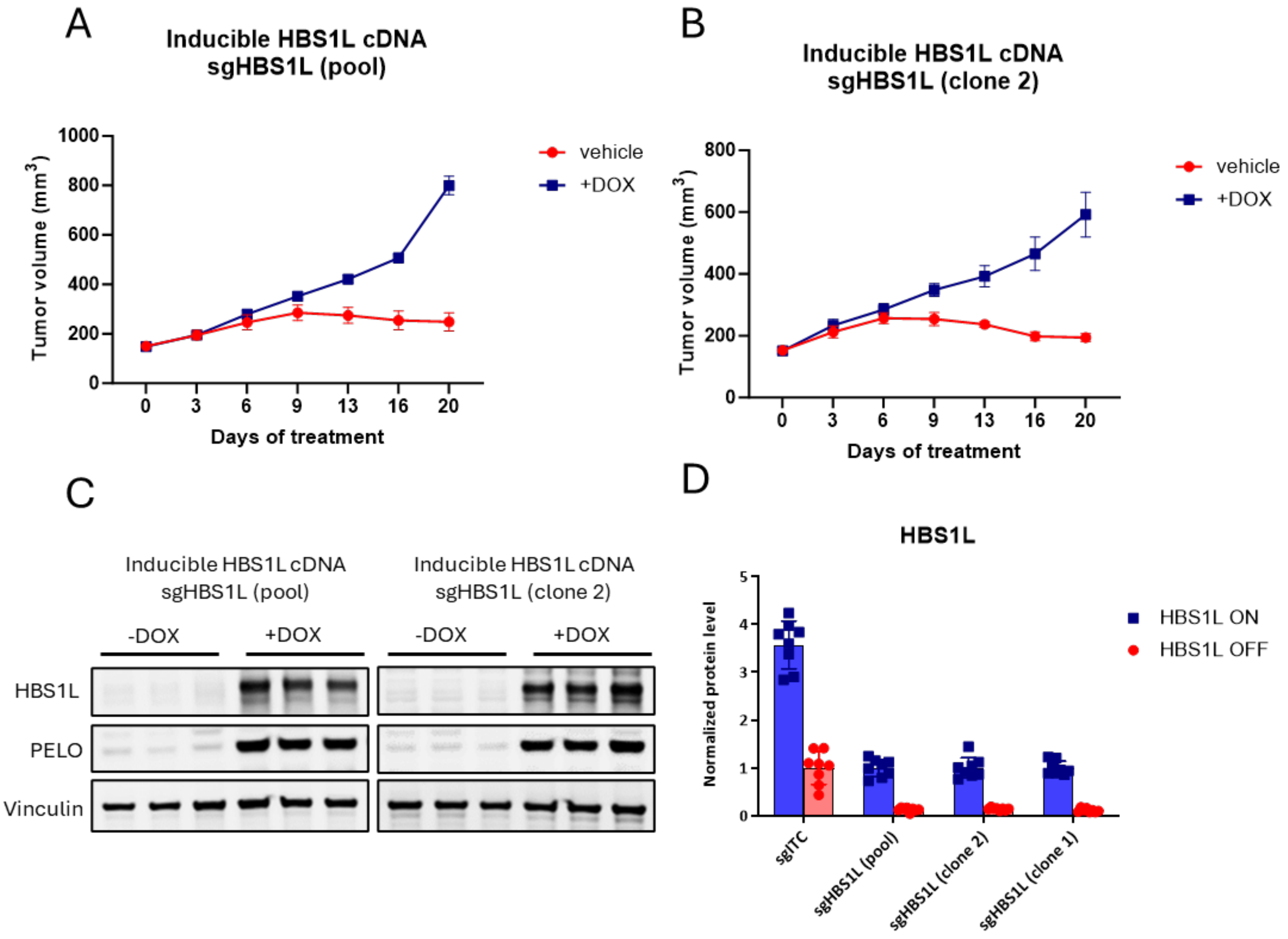

**Supplementary Figure S6. HBS1L loss eliminates tumor growth in CDX model of FOCAD-deleted pancreatic cancer in pooled and single cell clone format**

A, B) Growth of tumors in mice (N=8 animals/arm) implanted with MIAPACA2 cells expressing a doxycycline-inducible cDNA encoding HBS1L cDNA and knockout of endogenous HBS1L. Cells were implanted as a pooled population (A) or as a single cell clone with complete frameshift mutation in the endogenous *HBS1L* gene as confirmed by Sanger sequencing (B). Mice were fed chow with (+DOX) or without (vehicle) doxycycline to maintain or withdraw, respectively, expression of HBS1L. Data represent N=8 animals/arm, error bars represent ±

SEM. C, D) Representative image of HBS1L and PELO levels (C) and quantification (D) of HBS1L level detected by Western blotting of tumors at endpoint. Quantification represents mean of N=8 animals/arm, error bars represent ± SD.

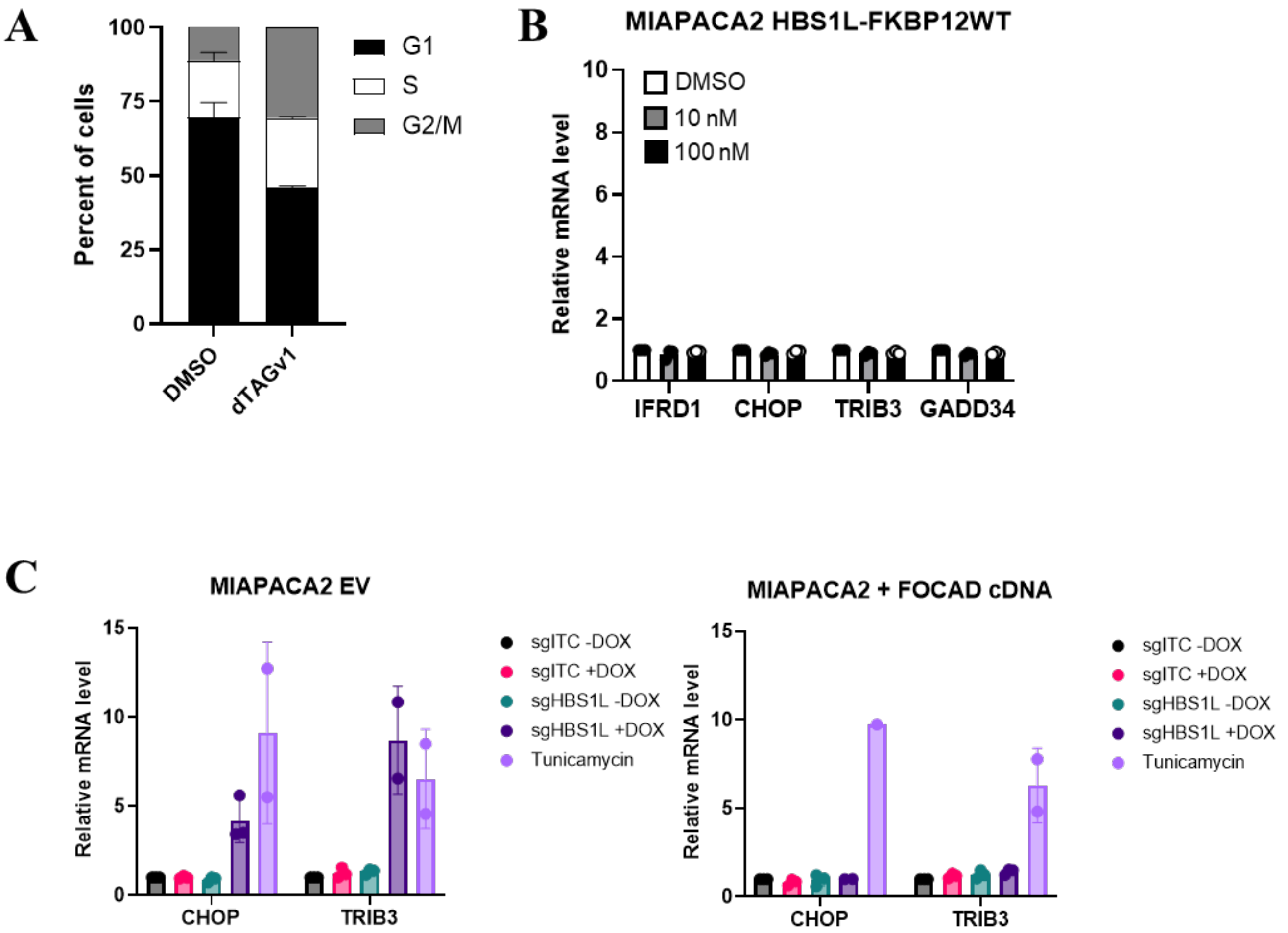

**Supplementary Figure S7. Degradation of HBS1L leads to cell cycle arrest in cells expressing an HBS1L-dTAG fusion protein, and inactivation of HBS1L leads to the induction of a transcriptional cellular stress response specifically in the FOCAD-deleted context.**

A) Cell cycle analysis of MIAPACA2 cells expressing a dTAG-fused *HBS1L* cDNA covering endogenous *HBS1L* knockout and treated with 0.11 µM dTAGv1 to induce degradation of HBS1L. Data represent the mean of 2 biological replicates, error bars represent ± SD. B) MIAPACA2 cells expressing an *HBS1L* cDNA fused to the non-degradable FKBP12WT control degron and with endogenous *HBS1L* knockout were treated with dTAGv1 for 24 hours, then samples were evaluated for the indicated transcripts by qPCR. Data represent the mean of 3 biological replicates, error bars represent ±SD. C) MIAPACA2 cells that lack (EV) or have (FOCAD cDNA) FOCAD reconstitution and express a DOX-inducible *HBS1L* cDNA covering endogenous *HBS1L* knockout (sgHBS1L) or an intron-cutting control sgRNA (sgITC). Upon DOX withdrawal, exogenous *HBS1L* expression is removed. Tunicamycin treatment (2 μg/mL, 24h) is used as a positive control for induction of the cellular stress response. Data represent the mean of 2 biological replicates, error bars represent ± SD.

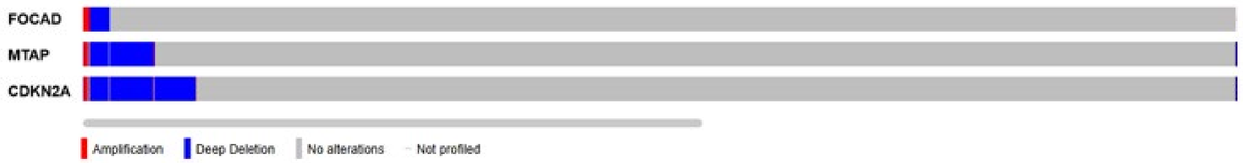

**Supplementary Figure S9. *FOCAD*, *MTAP*, and *CDKN2A* are frequently co-deleted in patient tumor samples.**

cBioportal (cBioportal.org) analysis of a curated set of non-redundant studies reporting copy number variation in *FOCAD*, *MTAP*, or *CDKN2A*. Samples that were not profiled for all three genes were not included.
